# Oxidative stress sensing by the translation elongation machinery promotes production of detoxifying selenoproteins

**DOI:** 10.1101/2025.10.13.682107

**Authors:** Frederick Rehfeld, Claire Lundstrom, He Zhang, Joshua T. Mendell

## Abstract

Selenocysteine, incorporated into polypeptides at recoded termination codons, plays an essential role in redox biology. Using GPX1 and GPX4, selenoenzymes that mitigate oxidative stress, as reporters, we performed genome-wide knockout screens to identify regulators of selenocysteine incorporation. This revealed that selenoprotein production is limited by ribosome collisions that occur at inefficiently decoded selenocysteine codons. Accordingly, slowed translation elongation reduced collisions and enhanced selenocysteine decoding. Oxidative stress also slowed translation elongation and augmented selenoprotein production. We identified translation elongation factor EEF1G as a sensor of oxidized glutathione that couples the cellular redox state to translation elongation rate. Oxidative stress sensing by EEF1G slows translation, enhancing production of detoxifying selenoproteins to restore homeostasis. These findings reveal how programmed ribosome collisions enable gene regulation in response to stress.

## Introduction

Signaling pathways that dynamically regulate mRNA translation play a central role in the acute response to various forms of cellular stress. This principle is exemplified by the integrated stress response (ISR), a pathway in which a series of stress-induced kinases phosphorylate the initiation factor eIF2α, which suppresses general protein synthesis while increasing translation of specific factors that help mitigate stress (*1*). In addition to the ISR, it has recently become clear that the protein synthesis machinery itself serves as a sentinel for cellular stress that can elicit potent signaling responses. Ribosomes are especially well suited to function as sensors of cellular stress due to their exceptional abundance and the highly dynamic nature of mRNA translation. Many insults, including oxidative stress and irradiation, can damage ribosomes and mRNAs, resulting in ribosomal stalling and collisions (*2–5*). The collided ribosome interface is now recognized as a platform for assembly of complexes that initiate multiple stress-induced signaling cascades (*6, 7*). The scope of ribosome collision events occurring within a cell is directly proportional to the signaling response. Local ribosome stalling and collisions on an individual mRNA lead to inhibition of translation initiation and decay of the transcript *in cis* (*8, 9*). Widespread collisions, however, activate the ISR kinase GCN2, resulting in global initiation shutdown, and can further activate the ribotoxic stress response (RSR) through the kinase ZAKα (also known as MAP3K20), which can induce cell cycle arrest or apoptosis, depending on the magnitude of stress (*5, 7, 10, 11*). These signaling events position translating ribosomes as key nodes in cellular stress sensing and response pathways.

Efficient translation, which helps mitigate ribosome collisions in unstressed cells, is promoted by the activity of translational GTPases (trGTPases) that serve as initiation, elongation, and termination factors (*12*). During canonical translation elongation, rapid decoding is facilitated by the trGTPase EEF1A which delivers aminoacyl-tRNAs. Similarly, termination is mediated by the decoding factor eRF1 in complex with the trGTPase eRF3. The activity of these factors ensures that the rate of miscoding or stop codon readthrough is generally low (<10^-4^ per codon) (*13*).

However, in specific contexts, targeted recoding of termination codons to non-canonical amino acids can occur. While pervasive in all three domains of life, this mechanism is specifically used in eukaryotes to generate proteins containing selenocysteine, a non-canonical amino acid that plays a key role in metazoan redox biology.

Selenocysteine is incorporated into nascent peptides through recoding of UGA termination codons (*14*). In eukaryotes, selenocysteine decoding is specified by the presence of a selenocysteine insertion sequence (SECIS), located in the 3’ untranslated region (UTR) of selenoprotein-encoding transcripts (*15*). SECIS is bound by the RNA-binding protein SECISBP2 (*16, 17*), which recruits the selenocysteine-specific elongation factor EEFSEC together with the selenocysteine-tRNA (tRNA^SEC^) to the ribosome (*18–20*). These selenocysteine-decoding ribosome complexes are known as selenosomes (*18*) (**Fig. 1A**). The selenocysteine pathway differs from canonical decoding in several ways. tRNA^SEC^ is the largest known tRNA (∼90nt) (*21*) and possesses unique structural features that distinguish it from canonical tRNAs, enabling its exclusive recognition by EEFSEC. Furthermore, tRNA^SEC^ undergoes a unique charging mechanism involving aminoacylation by serine-tRNA ligase (SARS1) followed by addition of selenium to the serine-charged tRNA following a phosphorylation step (*22, 23*).

**Figure 1.**
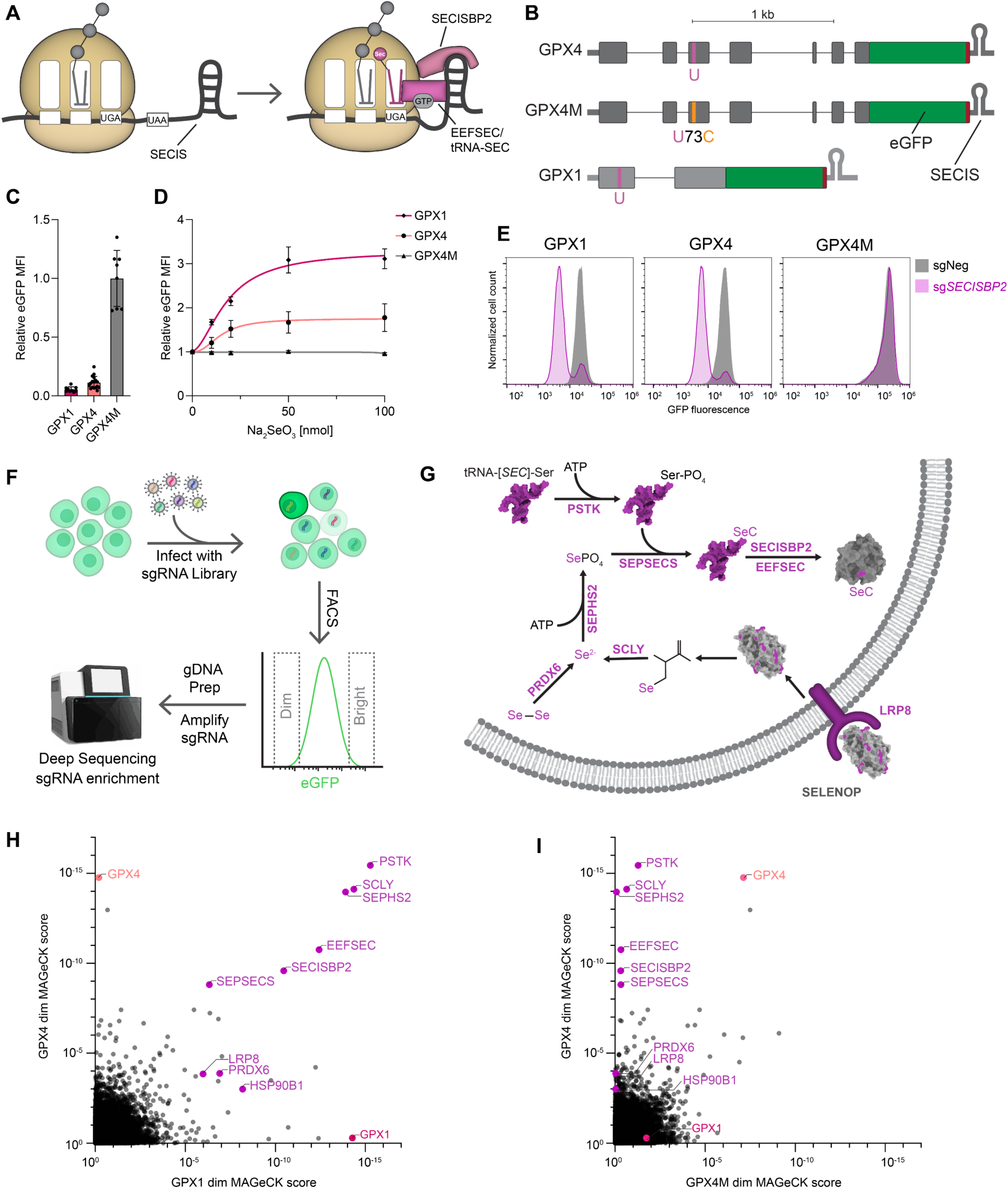
Genome-wide CRISPR-Cas9 knockout screens identify regulators of selenocysteine decoding. (**A**) Schematic representation of selenocysteine decoding. (**B**) GPX4 and GPX1 selenocysteine decoding reporter design. U, selenocysteine; C, cysteine. (**C**) Fluorescence of GPX1, GPX4, and GPX4M HCT116 reporter clones. Mean fluorescence intensity (MFI) was quantified by flow cytometry and normalized to GPX4M. Mean ± SD shown, with individual clones plotted. n=12 for GPX1, n=17 for GPX4, n=10 for GPX4M (biological replicates). (**D**) Sodium selenite dose response of GPX reporters. MFI was normalized to 0 nmol condition. Mean ± SD with four parameter logistic fit plotted. n=3 biological replicates each for GPX1, GPX4, and GPX4M at all tested doses. (**E**) Flow cytometry histograms of reporter cell lines transduced with lentiviral constructs encoding Cas9 and either negative control sgRNA (sgNeg) or sgRNA targeting *SECISBP2*. (**F**) Schematic of CRISPR-Cas9 knockout screen. Following transduction of HCT116 reporter clones with a pooled lentiviral CRISPR library, the brightest and dimmest cells were collected by FACS, followed by analysis of sgRNA enrichment in sorted populations by high-throughput sequencing. (**G**) Known factors required for the uptake, biosynthesis and incorporation of selenocysteine. (**H**) and (**I**) Results of the dim screen for factors required for selenocysteine decoding. MAGeCK gene enrichment scores plotted for GPX4 *vs.* GPX1 reporters (H) or GPX4 *vs.* GPX4M reporters (I). Known genes in the selenocysteine pathway are labeled in purple.

The evolutionary benefit of maintaining such an intricate and energy intensive decoding pathway is thought to lie in the unique chemical properties of selenocysteine. Its lower pK_a_ and higher nucleophilicity make selenocysteine more reactive than cysteine and therefore critical for catalysis by enzymes that mitigate oxidative stress and maintain cellular redox homeostasis (*24*). This is illustrated by the fact that the majority of the 25 human selenoproteins harbor oxidoreductase domains. Among the most important selenoenzymes is GPX4, an essential glutathione-dependent phospholipid peroxidase. Loss of either the selenocysteine decoding machinery (*25*) or GPX4 (*26*) results in early embryonic lethality in mice. GPX4 protects cells from the iron-dependent accumulation of phospholipid peroxides (PLOOH), thereby preventing a form of cell death known as ferroptosis (*27*). Cancer cells have been shown to be especially vulnerable to lipid peroxidation and ferroptosis (*28–30*), thus making GPX4, and the selenocysteine decoding pathway in general, attractive targets for drug development (*31*).

Selenocysteine decoding is significantly less efficient than canonical A-site decoding (*32*), raising the possibility that this pathway is subject to currently unknown regulatory mechanisms. Nevertheless, despite the importance of GPX4 and other selenoproteins in mammalian physiology and disease, regulators of selenocysteine incorporation beyond the core decoding machinery have yet to be identified. To comprehensively identify regulators of this pathway, we performed genome-wide CRISPR-Cas9 knockout screens leveraging glutathione peroxidases *GPX4* and *GPX1* as reporters of selenocysteine decoding. These screens successfully recovered all previously known factors required for the biosynthesis and incorporation of selenocysteine. Furthermore, we found that, due to the inefficient decoding of selenocysteine codons, selenoprotein translation is limited by ribosome collisions that basally occur on these mRNAs. A global slowdown in translation elongation reduced these collisions, resulting in a marked increase in selenoprotein production. Intriguingly, we found that oxidative stress also slowed translation elongation and similarly increased selenocysteine incorporation. We identified the eukaryotic elongation factor 1 (EEF1) subunit EEF1G as a sensor that directly couples the cellular redox status to elongation speed. These findings uncovered a novel regulatory circuit in which direct regulation of translation elongation by oxidative stress promotes the production of detoxifying selenoproteins, thereby contributing to maintenance of cellular redox homeostasis.

## Results

### Genome-wide CRISPR-Cas9 knockout screens identify regulators of selenocysteine decoding

To monitor the efficiency of selenocysteine decoding, we generated reporter constructs based upon the essential glutathione peroxidase and selenoenzyme *GPX4*. The complete *GPX4* gene, including introns and UTRs, was cloned into a constitutive mammalian expression vector and the coding sequence (CDS) of enhanced green fluorescent protein (eGFP) was fused to the 3’ end of the open reading frame (**Fig. 1B**). To distinguish effects on selenocysteine-decoding from other post-transcriptional regulatory mechanisms that impact *GPX4* expression, we generated a reporter construct in which the selenocysteine codon was mutated to a cysteine codon, referred to as GPX4 mutant (GPX4M). A third reporter construct based on another selenoenzyme, the cytosolic glutathione peroxidase GPX1, was also generated to enable identification of factors that generally impact selenocysteine decoding.

Using the PiggyBac transposase system (*33, 34*), we established stable clonal HCT116 cell lines expressing the reporter constructs. Flow cytometry of these reporter cell lines revealed that eGFP expression was approximately 10-to 20-fold lower in GPX4 and GPX1 reporter cells compared to GPX4M cells (**Fig. 1C**). This is consistent with results obtained using luciferase reporters of selenocysteine decoding (*32*) and reflects the relative inefficiency of selenocysteine incorporation compared to canonical amino acids. Supplementation of culture medium with inorganic selenium led to a dose-dependent increase in expression of GPX4 and GPX1 reporters without affecting the GPX4M reporter (**Fig. 1D**). Conversely, CRISPR-mediated loss of function of canonical selenocysteine decoding factors SECISBP2 or PSTK resulted in a marked decrease in GPX4 and GPX1, but not GPX4M, reporters (**Fig. 1E and fig. S1A**). These reporters therefore provided a fluorescent readout of the efficiency of selenocysteine incorporation.

We next leveraged this system to perform genome-wide loss-of-function screens to identify regulators of selenocysteine decoding. Two independent clonal cell lines expressing each reporter were infected at low multiplicity of infection with a lentiviral CRISPR-Cas9 single guide RNA (sgRNA) library targeting ∼19,000 human protein-coding genes (*35*). Fluorescence-activated cell sorting (FACS) was used to collect the brightest and dimmest cells, and enrichment of sgRNAs in sorted over unsorted pools was determined by high-throughput sequencing (**Fig. 1F**). We first examined the genes that were targeted by sgRNAs that were significantly enriched in the dim population, representing candidates that are required for selenocysteine incorporation (**table S1**). Indeed, all known factors required for biosynthesis of tRNA^SEC^ (*SCLY, SEPHS2, PSTK, SEPSECS*) and selenocysteine decoding (*SECISBP2, EEFSEC*) scored as highly significant hits in the GPX4 and GPX1 screens, but not in the GPX4M screen (**Fig. 1, G to I**). We also identified the genes encoding LRP8 (also named ApoER2), which is the main receptor for the selenium storage protein SELENOP (*36, 37*), the ER-resident LRP8 chaperone HSP90B1 (*38*), and PRDX6, a peroxiredoxin enzyme that was recently reported to promote selenium utilization by reducing cellular diselenides (*39–41*), as significant hits in the selenocysteine-dependent screens. CRISPR-mediated knockout of *LRP8, HSP90B1*, and *PRDX6* confirmed their requirement for expression of the selenocysteine-dependent reporters (**fig. S1, B to D**). Unbiased examination of hits using Gene set enrichment analysis (GSEA) revealed that pathways related to transcriptional regulation of the reporters and selenocysteine incorporation were the only significantly enriched categories shared between the GPX1 and GPX4 dim screens (**fig. S1, E and F**). These results demonstrated that this screening platform provided a sensitive approach for the identification of genes that regulate selenoprotein production.

### Negative regulators of selenocysteine decoding

Genes whose loss of function resulted in an increase in expression of the GPX reporters (‘bright screen’ hits) represent potential negative regulators of selenocysteine decoding (**table S2**). A notable example was thioredoxin reductase 1 (*TXNRD1*), a selenoenzyme and the main cytosolic reductase for the antioxidant thioredoxin system, which was the top hit in both selenocysteine-dependent bright screens but was not enriched in the GPX4M screen (**Fig. 2, A and B**). We validated that *TXNRD1* knockout increased selenocysteine-dependent reporter expression but did not affect the GPX4M reporter (**Fig. 2C**). GSEA was used to systematically examine pathways whose disruption resulted in de-repression of the selenocysteine-dependent GPX4 and GPX1 reporters. This revealed that the most significantly enriched gene sets largely fell into three categories: mitochondrial function, translation elongation, and ribosome biogenesis (**fig. S2, A to D**).

**Figure 2.**
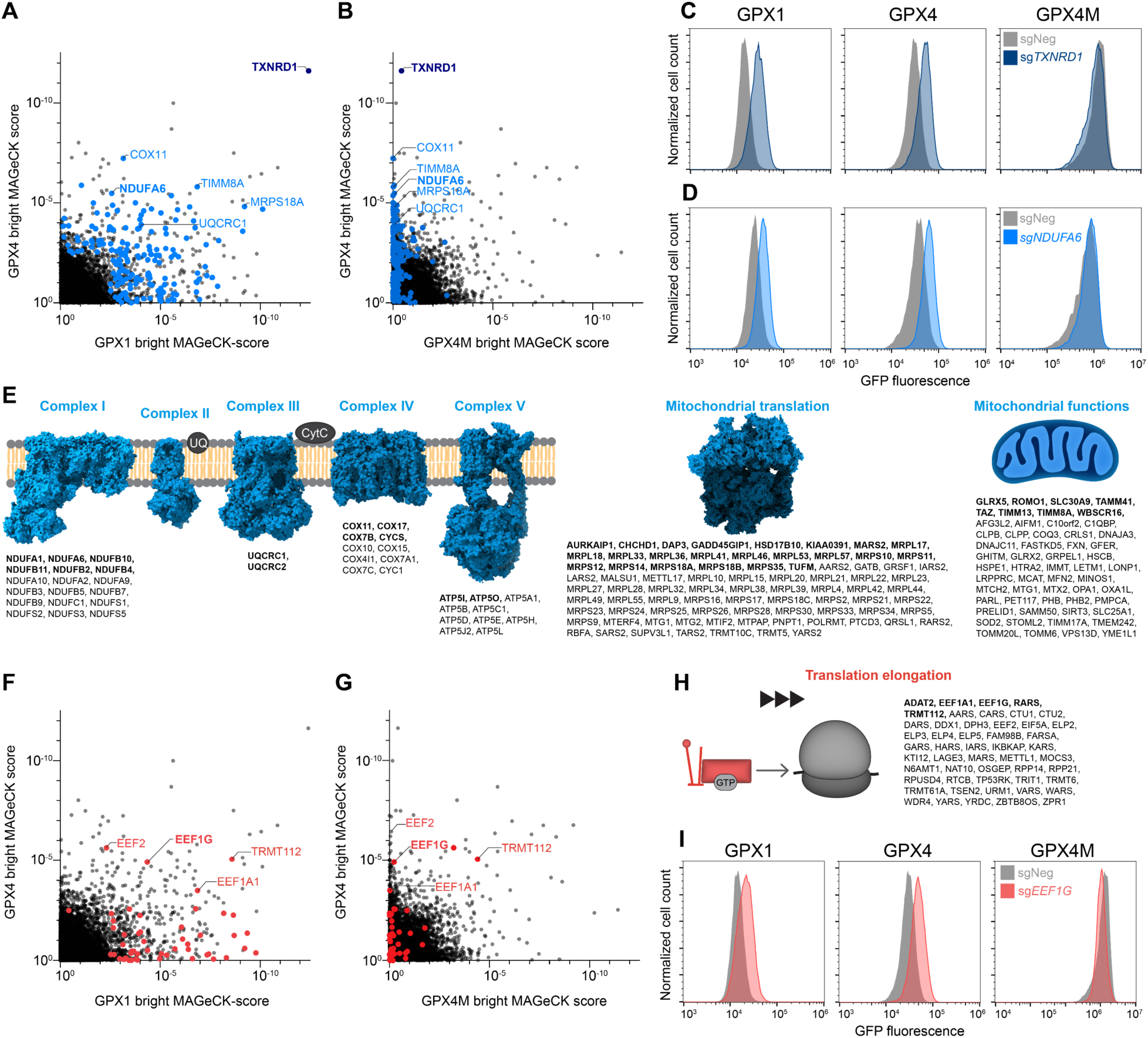
Negative regulators of selenocysteine decoding. (**A**) and (**B**) Results of the bright screen for negative regulators of selenocysteine decoding. MAGeCK gene enrichment scores plotted for GPX4 *vs.* GPX1 reporters (A) or GPX4 *vs.* GPX4M reporters (B). Significantly enriched genes in either the GPX1 or GPX4 screen (p<0.01) with roles related to mitochondrial function are labeled in blue. (**C**) and (**D**) Flow cytometry histograms of GPX reporter cell lines transduced with lentiviral constructs encoding Cas9 and the indicated sgRNAs. (**E**) Significantly enriched genes in either the GPX1 or GPX4 bright screen (p<0.01) categorized by mitochondrial function. Significant hits in both screens are labeled in bold. (**F**) and (**G**) Significantly enriched genes in either the GPX1 or GPX4 bright screen (p<0.01), plotted as in (A) and (B). Genes with functions related to translation elongation are labeled in red. (**H**) Significantly enriched genes in either the GPX1 or GPX4 bright screen (p<0.01) with functions related to translation elongation. Significant hits in both screens are labeled in bold. (**I**) Flow cytometry histograms of GPX reporter cell lines transduced with lentiviral constructs encoding Cas9 and the indicated sgRNAs.

Genes involved in mitochondrial function made up the largest category of genes retrieved in the selenocysteine-dependent bright screens. A cumulative total of 174 genes in this category were recovered as significant hits (p < 0.01) in the GPX4 and/or GPX1, but not GPX4M, screens (**Fig. 2, A, B, and E**). These genes encoded components of electron transport chain (ETC) complexes I, III, IV and V, as well as proteins involved in mitochondrial translation and many other mitochondrial functions. Indeed, knockout of NDUFA6, a component of ETC complex I, increased GPX1 and GPX4 reporter expression without affecting GPX4M (**Fig. 2D**). Many genes related to cytoplasmic translation elongation and ribosome biogenesis were also enriched in the selenocysteine-dependent screens (**Fig. 2, F to H, and fig. S3, A to C**). These genes included canonical elongation factors and other genes involved in amino acid decoding. We confirmed that loss of EEF1G, an essential EEF1 subunit, resulted in upregulation of the selenocysteine reporters, while modestly reducing the expression of the GPX4M reporter (**Fig. 2I**). Based on these findings, we set out to investigate how perturbations of translation and mitochondrial function could increase selenocysteine decoding.

### Inhibition of translation elongation enhances selenoprotein synthesis by reducing ribosome collisions

The results of the CRISPR screen suggested that inhibition of translation elongation paradoxically increases selenoprotein synthesis. Most genes in this category that were retrieved in the CRISPR screen play a role in canonical A-site decoding, including elongation factors as well as proteins that process, modify, or charge tRNAs (**Fig. 2H**). We expanded our validation of this category of genes by showing that CRISPR-mediated loss of the EEF1 complex subunits *EEF1A1* and *EEF1G*, *EEF2,* and tRNA modifying (*ELP5*) and charging (*VARS*) factors led to an upregulation of selenocysteine reporter expression (**Fig. 2I**, **Fig. 3A, and fig. S3, D to F**).

**Figure 3.**
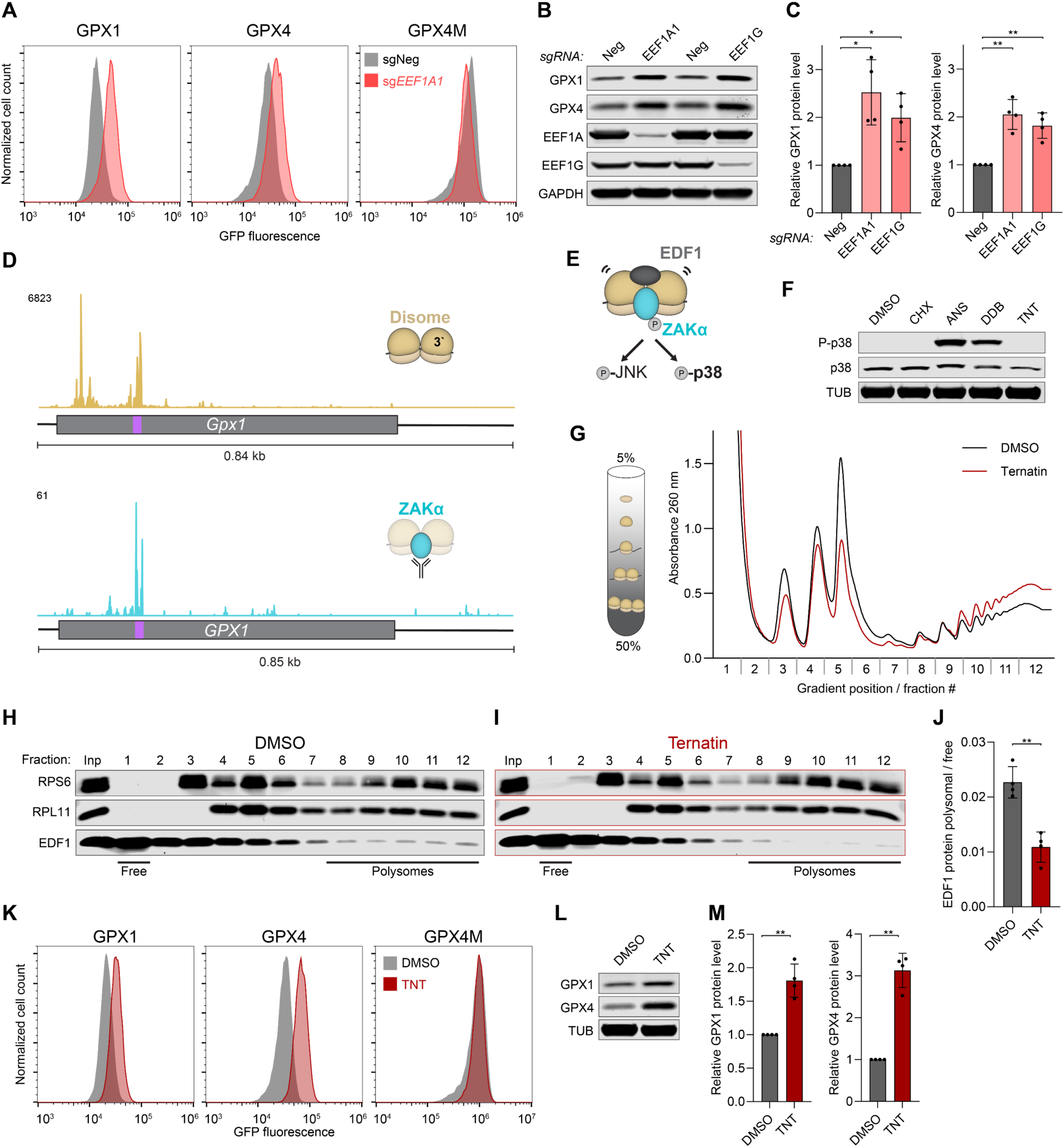
Slowed translation elongation enhances selenocysteine decoding by reducing ribosome collisions. (**A**) Flow cytometry histograms of GPX reporter cell lines transduced with lentiviral constructs encoding Cas9 and the indicated sgRNAs. (**B**) and (**C**) Western blot analysis (B) and associated quantification (C) of GPX1 and GPX4 protein levels in HCT116 cells following lentiviral delivery of Cas9 and the indicated sgRNAs. Protein abundance was normalized to GAPDH and to sgNeg. Mean ± SD plotted. n=4 biological replicates. *P* values were calculated by one-sample t-test. *p < 0.05, **p < 0.01. (**D**) Disome profiling (*45*) (upper) and ZAKα selective ribosome profiling (*7*) (lower) read coverage on mouse *Gpx1* and human *GPX1,* respectively. For disomes, reads were plotted based on position of the P-site of the leading ribosome. Coding sequences indicated with grey boxes, selenocysteine codons indicated in purple. (**E**) Activation of the ribotoxic stress response. Recruitment of ZAKα to collided ribosomes leads to phosphorylation of p38 and JNK and associated downstream signaling. EDF1 is also recruited to the collided ribosome interface. (**F**) Western blot analysis of p38 phosphorylation in HEK293T cells following treatment for 7 minutes with DMSO, 0.5 µg/ml cycloheximide (CHX), 0.5 µg/ml anisomycin (ANS), 0.75 µM didenmin B (DDB), or 0.75 µM ternatin (TNT). (**G**) Sucrose gradient sedimentation of ribosomes. Cells were treated with DMSO or 0.75 µM TNT for 7 minutes prior to harvesting. (**H**) and (**I**) Western blot analysis of collected sucrose gradient fractions after treatment with DMSO (H) or 0.75 µM ternatin (I). Inp, input prior to sedimentation. (**J**) Ratio of polysome associated EDF1 (fractions 8-12) over cytosolic EDF1 (fraction 1) quantified by western blot. Mean ± SD plotted. n=4 biological replicates. *P* value was calculated by student’s t-test. **p < 0.01. (**K**) Flow cytometry histograms of GPX reporter cell lines treated with DMSO or 0.5 µM TNT for 48 hours. (**L**) and (**M**) Western blot analysis (L) and associated quantification (M) of GPX1 and GPX4 protein levels in HCT116 cells following treatment with either DMSO or 0.5 µM TNT for 72 hours. Protein abundance was normalized to GAPDH and to DMSO-treated condition. Mean ± SD plotted. n=4 biological replicates. *P* values were calculated by one-sample t-test. **p < 0.01.

Conversely, the GPX4M reporter was unaffected or decreased upon loss of these genes. Endogenous GPX1 and GPX4 protein abundance was similarly elevated upon EEF1A1 or EEF1G loss compared to control cells (**Fig. 3, B and C**).

Interestingly, we noticed that genes encoding ribosome rescue factors that clear stalled ribosomes from transcripts (*HBS1L, PELO, ABCE1*) (*42–44*) were also identified as significant hits in the GPX1 and GPX4 bright screens (**fig. S4, A to C**). We reasoned that due to inefficient decoding of selenocysteine codons, a subset of selenosomes might be recognized as stalled ribosomes and therefore cleared by ribosome rescue pathways. We confirmed that loss of *PELO* increased expression of the selenocysteine reporters (**fig. S4D**).

Given these results, we hypothesized that ribosomes stalled at selenocysteine codons might also be subject to collisions with trailing ribosomes. Indeed, disome peaks, indicative of ribosome collisions, were detected at a subset of selenocysteine codons in a previously-reported disome profiling dataset from mouse liver (*45*). Further analysis of these data confirmed the presence of a disome peak at the selenocysteine codon in *Gpx4* (**fig. S5A**), and revealed an even more robust peak at the *Gpx1* selenocysteine codon (**Fig. 3D**). To determine whether these disomes trigger signaling events associated with ribosome collisions, we examined ZAKα selective ribosome profiling data from HEK293T cells (*7*). ZAKα is a kinase that specifically associates with collided ribosomes. This binding induces ZAKα autophosphorylation which, in turn, activates the ribotoxic stress response (RSR) through phosphorylation of MAPKs p38 (MAPK14) and JNK (MAPK8) (*6, 7, 10, 11*) (**Fig. 3E**). A strong ZAKα peak was evident at the *GPX1* selenocysteine codon (**Fig. 3D**), while a discernable ZAKα peak was not detected on *GPX4* in this dataset, possibly due to lower transcriptome coverage provided by selective ribosome profiling (**fig. S5B**). Importantly, several other selenoprotein encoding transcripts (*SELENOF, SELENOH, SELENOM, SELENOO, SELENOS, SELENOW and TXNRD1*) exhibited robust disome peaks, and in most cases corresponding ZAKα peaks, at their respective selenocysteine codons (**fig. S5, A and B**). Thus, ribosome collisions that recruit ZAKα frequently occur during selenocysteine decoding.

The finding that selenocysteine codons trigger ribosome collisions provides a mechanism to explain how slowing of translation elongation increases selenoprotein production. Because ribosome collisions lead to ribosome disassembly and clearance of nascent peptides, while also decreasing translation initiation *in cis* (*8, 9, 46–48*), these collision events are expected to limit selenoprotein synthesis. Globally reduced translation elongation would reduce collisions by providing a window for completion of selenocysteine incorporation before a trailing ribosome reaches the selenosome. To directly test whether slowing of translation elongation reduces collisions and enhances selenocysteine decoding, we sought to recapitulate these effects using pharmacologic inhibition of translation elongation. A panel of translation inhibitors was tested to identify a compound that globally reduces ribosome collisions, as monitored by ZAKα-mediated p38 phosphorylation and polysome association of the collision marker EDF1 (*49*). As previously reported, treatment with the peptidyl-transferase inhibitor anisomycin (ANS) (*6, 7*) or the potent EEF1A inhibitor didemnin B (DDB) (*50*) strongly induced ribosome collisions, as indicated by p38 phosphorylation (**Fig. 3, E and F**) and increased polysome association of EDF1 (**fig. S6, A to C**). On the other hand, treatment with the translocation inhibitor cycloheximide (CHX) (*51*) did not induce p38 phosphorylation and had little effect on polysome association of EDF1 (**Fig. 3F and fig. S6D**). Most notably, the weak EEF1A inhibitor ternatin (TNT) (*52, 53*) uniquely reduced the global level of ribosome collisions, as indicated by a lack of p38 phosphorylation and lower levels of EDF1 on polysomes compared to control-treated cells (**Fig. 3, F to J**). As predicted, this reduced burden of collisions in TNT-treated cells was associated with increased expression of the selenocysteine-dependent GPX1 and GPX4 reporters, but not the GPX4M construct (**Fig. 3K**). Endogenous GPX1 and GPX4 protein abundance also increased upon TNT treatment (**Fig. 3, L and M**). Expanding these analyses beyond GPX1 and GPX4, we observed that CRISPR-mediated knockout of *EEF1A1* or *EEF1G*, or TNT treatment, similarly increased expression of SELENOF and SELENOH, encoded by transcripts that exhibited strong collision signatures in the disome and ZAKα selective ribosome profiling (**fig. S5, A and B, and fig. S6, E and F**).

These data demonstrate that selenoprotein production is limited by ribosome collisions which occur at slowly-decoded selenocysteine codons, an effect that can be mitigated by a global slowing of translation elongation. We note that a similar mechanism likely explains our finding that depletion of factors required for ribosome biogenesis increased selenocysteine decoding, since reduced ribosome load would be expected to reduce collisions as well.

### Oxidative stress and redox imbalance increase selenoprotein synthesis

Components of the ETC and other genes required for proper mitochondrial function represented the largest class of hits whose loss increased expression of the selenocysteine-dependent reporters (**Fig. 2E**). Accordingly, genetic loss or pharmacological inhibition of subunits from complexes I, III or V, or loss of mitochondrial ribosomal protein-encoding genes, resulted in increased GPX1 and GPX4, but not GPX4M, reporter expression (**Fig. 2D**, **Fig. 4, A to D, and fig. S7, A to D**). Perturbation of oxidative phosphorylation is known to cause mitochondrial electron leakage and a concomitant increase in cellular reactive oxygen species (ROS) (*54, 55*). We documented that this expected increase in cellular ROS resulted from depletion of components of complexes I, III and V, but did not occur after loss of TXNRD1, the top hit in the bright screens and the key enzyme in the antioxidant thioredoxin pathway (**Fig. 2, A and B, Fig. 4E, and fig. S7E**). However, all of these perturbations decreased the ratio of reduced (GSH) to oxidized (GSSG) glutathione, indicative of a more oxidizing cellular environment (**fig. S7F**). We were intrigued by these findings since increased production of detoxifying selenoproteins during conditions of oxidative stress would allow for immediate and homeostatic stress mitigation.

**Figure 4.**
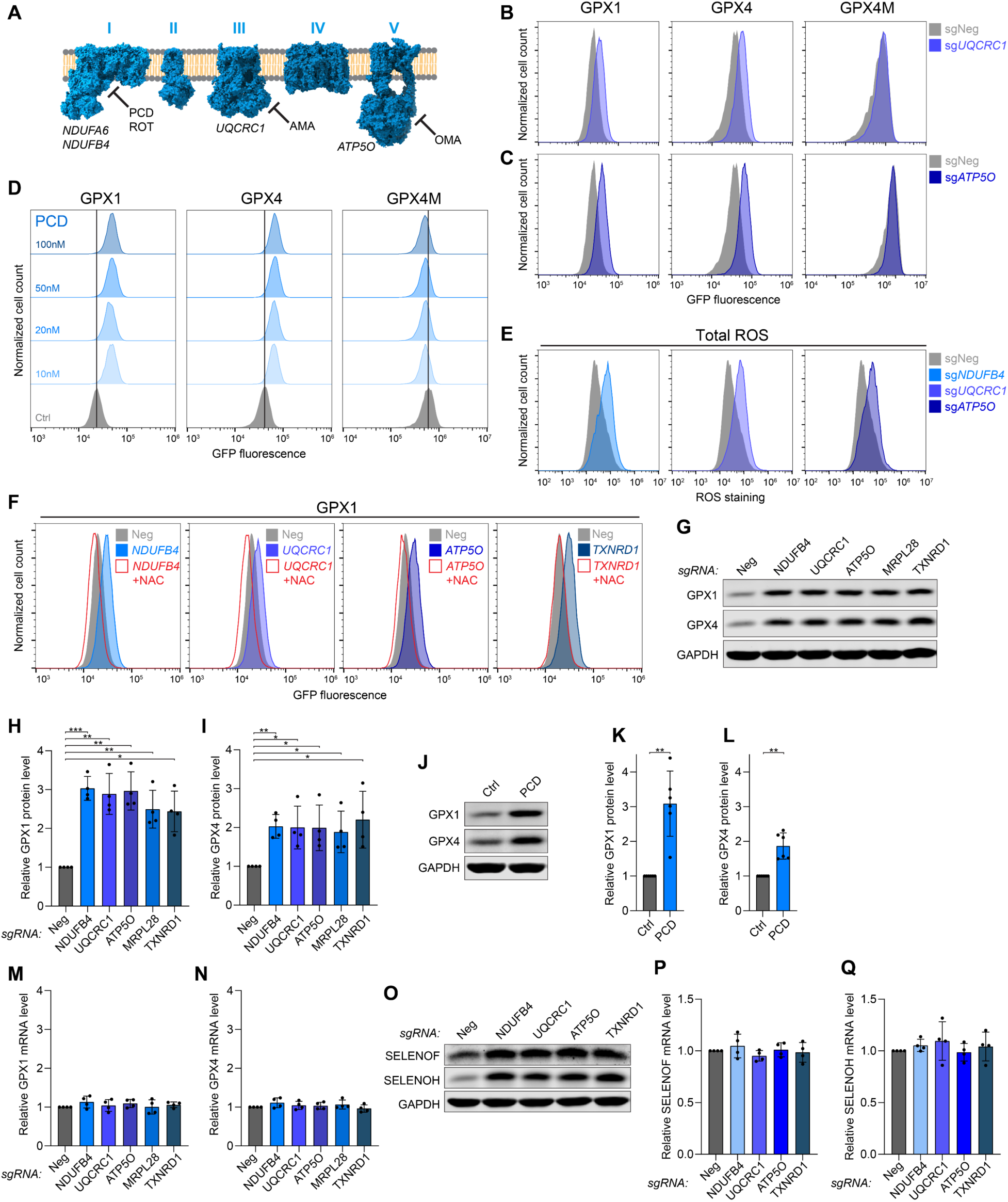
Oxidative stress increases selenocysteine decoding. (**A**) Schematic of mitochondrial electron transport chain showing targets of inhibitors and representative genes selected for loss-of-function studies. PCD, piericidin; ROT, rotenone; AMA, antimycin A; OMA, oligomycin A. (**B**) and (**C**) Flow cytometry histograms of GPX reporter cell lines transduced with lentiviral constructs encoding Cas9 and the indicated sgRNAs. (**D**) Flow cytometry histograms of GPX reporter cell lines treated with increasing doses of PCD for 72 hours. (**E**) Total cellular ROS quantification by flow cytometry in HCT116 cells following lentiviral delivery of Cas9 and the indicated sgRNAs. (**F**) Flow cytometry histograms of GPX1 reporter cells transduced with lentiviral constructs encoding Cas9 and the indicated sgRNAs with or without treatment with N-acetyl cysteine (NAC). (**G**) to (**I**) Western blot analysis (G) and associated quantification (H and I) of GPX1 and GPX4 protein levels in HCT116 cells following lentiviral delivery of Cas9 and the indicated sgRNAs. Protein abundance was normalized to GAPDH and to sgNeg. Mean ± SD plotted. n=4 biological replicates. *P* values were calculated by one-sample t-test. *p < 0.05, **p < 0.01, ***p < 0.001. (**J**) to (**L**) Western blot analysis (J) and associated quantification (K and L) of GPX1 and GPX4 protein levels in HCT116 cells treated with DMSO or 20 nM PCD for 72 hours. Protein abundance was normalized to GAPDH and to mock-treated sample. Mean ± SD plotted. n=6 biological replicates. *P* values were calculated by one-sample t-test. *p < 0.05, **p < 0.01. (**M**) and (**N**) qRT-PCR analysis of *GPX1* (M) and *GPX4* (N) transcript abundance in HCT116 cells transduced with lentiviral constructs expressing Cas9 and the indicated sgRNAs. Expression was normalized to housekeeping gene *Oaz1* and negative control sgRNA. Mean ± SD plotted. n=4 biological replicates. (**O**) Western blot analysis of SELENOF and SELENOH in HCT116 cells following lentiviral delivery of Cas9 and the indicated sgRNAs. (**P**) and (**Q**) qRT-PCR analysis of *SELENOF* (M) and *SELENOH* (N) transcript abundance in HCT116 cells transduced with lentiviral constructs expressing Cas9 and the indicated sgRNAs. Expression was normalized to housekeeping gene *Oaz1* and negative control sgRNA. Mean ± SD plotted. n=4 biological replicates.

Moreover, it was previously reported that exogenously induced oxidative stress increases the synthesis of selenoproteins (*56*), although the mechanism underlying this effect is not known.

To test if ROS and the altered cellular redox state were indeed responsible for upregulation of selenocysteine reporters upon loss of mitochondrial proteins or TXNRD1, knockout cells were treated with N-acetyl cysteine (NAC), an antioxidant and potent reducing agent that can restore redox homeostasis by reducing cellular dithiols (*57*). Treatment of cells with NAC completely reversed GPX1 reporter upregulation upon knockout of ETC components or *TXNRD1*, confirming the key role of oxidative stress in selenocysteine reporter upregulation under these conditions (**Fig. 4F**).

Consistent with the behavior of the reporters, endogenous GPX1 and GPX4 protein levels were induced by the loss of ETC components, the mitochondrial ribosomal protein MRPL28, or TXNRD1 (**Fig. 4, G to I**). Concordant effects were observed after treatment with the complex I inhibitor piericidin (PCD) (**Fig. 4, J to L**). Intriguingly, these perturbations did not increase *GPX1 or GPX4* transcript levels (**Fig. 4, M and N**). *SELENOF* and *SELENOH* were also induced at the protein, but not mRNA, level under these conditions (**Fig. 4, O to Q**), while *NQO1*, an NRF2 target gene that is transcriptionally upregulated upon oxidative stress (*58*), showed the expected mRNA induction (**fig. S7G**). These results, coupled with the observation that the GPXM reporter was not induced by these perturbations, suggested that oxidative stress increases selenoprotein abundance through a selenocysteine-dependent, post-transcriptional mechanism. To investigate whether post-translational regulation of GPX4 protein levels contributed to the observed increase in selenoprotein expression, we generated a reporter with a self-cleavable PT2A peptide sequence inserted between GPX4 and eGFP (**fig. S7H**). Since GPX4 and eGFP are co-translationally separated in cells expressing this reporter, any post-translational effects on GPX4 protein should not affect eGFP levels, but eGFP production will still be impacted by the efficiency of selenocysteine decoding. ETC or TXNRD1 loss of function strongly increased expression of this reporter, pointing to an effect on selenocysteine decoding (**fig. S7I**).

Altogether, these data provide strong evidence that selenocysteine decoding is stimulated by an oxidative cellular environment, explaining why disruption of a broad array of mitochondrial functions or the thioredoxin system induces selenoprotein production.

### Oxidative stress increases selenocysteine decoding by slowing translation elongation

To investigate the mechanism by which selenocysteine decoding is enhanced under oxidative conditions, we first examined the levels of rate limiting factors in the selenocysteine pathway upon oxidative stress. However, the abundance of key factors such as PSTK, SECISBP2, or EEFSEC were not increased following treatment with the complex I inhibitor PCD (**fig. S8A**).

Since it has been shown that oxidative stress inhibits translation elongation in yeast (*59, 60*), we next considered the possibility that oxidative stress may have similar effects in mammalian cells. As we demonstrated above, slowed elongation would reduce ribosome collisions at selenocysteine codons and thereby increase production of selenoproteins.

To test whether oxidative stress inhibits translation elongation in mammalian cells, we measured ribosome runoff in HCT116 cells by acutely blocking translation initiation with silvestrol (SIL) and assessing polysome levels over time. This revealed that while the overall abundance of polysomes was reduced upon genetic loss or pharmacological inhibition of ETC subunits, likely reflecting reduced translation initiation due to ISR induction, polysome runoff was also dramatically slowed (**Fig. 5, A and B, and fig. S8, B and C**). Similar results were obtained following treatment with the ROS inducing agent paraquat (**fig. S8, D and E**). Runoff assays performed by quantifying nascent peptides over time by puromycylation (*61*) similarly documented decelerated ribosome runoff in PCD treated cells (**fig. S8, F and G**).

**Figure 5.**
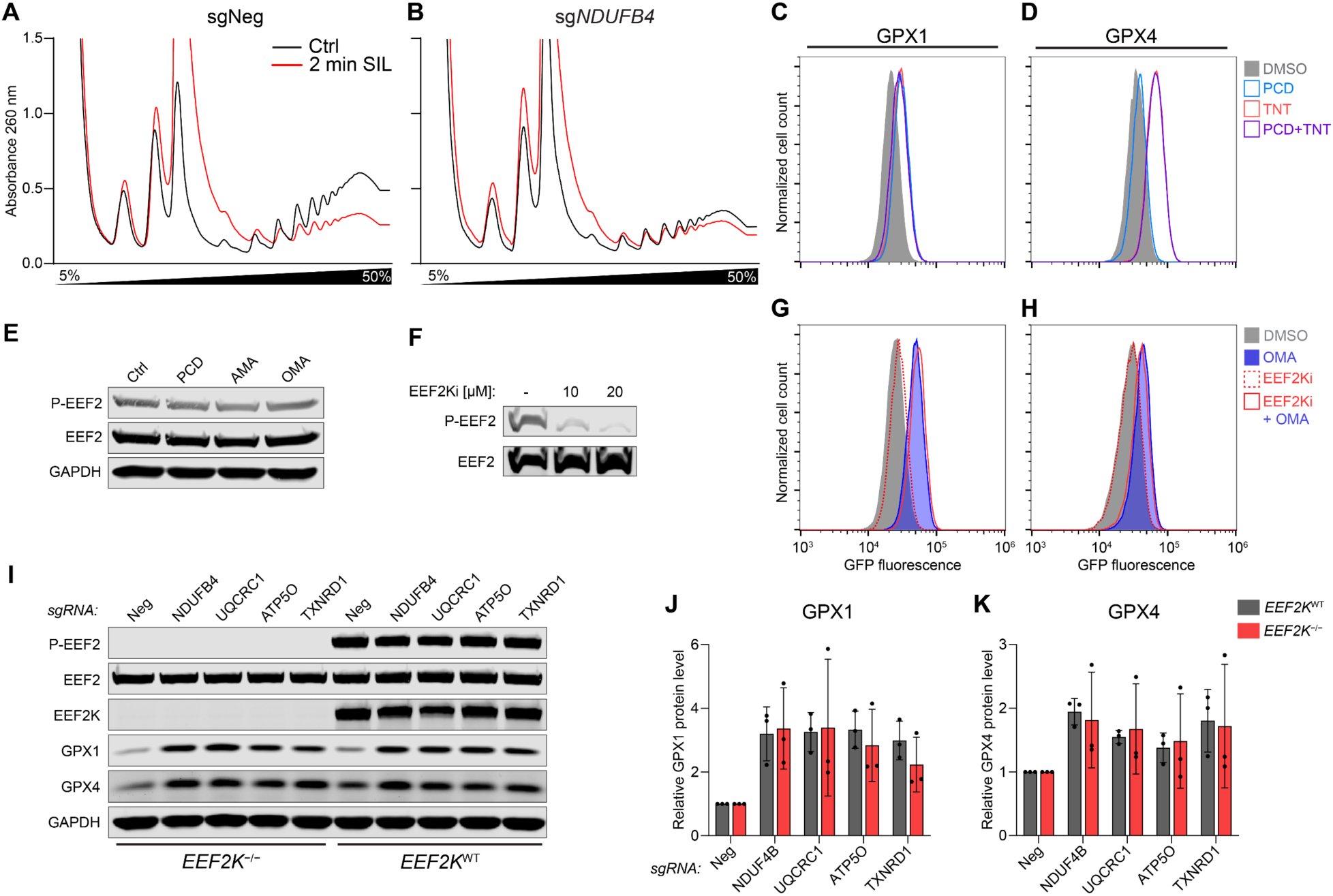
Oxidative stress slows translation elongation in an EEF2K-independent manner. (**A**) and (**B**) Ribosome runoff experiments in HCT116 cells transduced with lentiviral constructs expressing Cas9 and the indicated sgRNAs. Polysome abundance was monitored by sucrose gradient sedimentation before and after 2 minutes of treatment with the translation initiation inhibitor silvestrol (SIL; 2 µM). (**C**) and (**D**) Flow cytometry histograms of GPX reporter cells either untreated (Ctrl) or treated with 10 nM PCD, 0.75 µM TNT, or both PCD and TNT. (**E**) and (**F**) Western blot analysis of EEF2 phosphorylation in HCT116 cells with or without 24 hours of treatment with 20 nM of ETC inhibitors (E) or after treatment with the indicated doses of EEF2K inhibitor A-484954 (EEF2Ki) (F). (**G**) and (**H**) Flow cytometry histograms of GPX1 (G) and GPX4 (H) reporter cell lines treated with vehicle (DMSO) or the indicated compounds. (**I**) to (**K**) Western blot analysis (I) and quantification of GPX1 and GPX4 protein levels (J and K) in *EEF2K* wild type or knockout HCT116 cells transduced with lentiviral constructs encoding Cas9 and the indicated sgRNAs. Protein abundance was normalized to GAPDH and to sgNeg. Mean ± SD plotted. n=3 biological replicates.

Having established that oxidative stress reduces translation elongation speed in mammalian cells, we next performed epistasis experiments in which we treated selenocysteine reporter cells with either PCD (oxidative stress), TNT (slowed elongation), or both compounds simultaneously. If PCD and TNT impact selenocysteine decoding through independent pathways, we would expect that simultaneous treatment with both drugs would yield an additive phenotype.

However, GPX1 reporter expression was similarly increased upon treatment with PCD, TNT, or the combination (**Fig. 5C**). GPX4 expression was more strongly upregulated by TNT compared to PCD but showed no further increase upon co-treatment (**Fig. 5D**). These results suggested that oxidative stress increases selenocysteine decoding by slowing canonical translation elongation, providing evidence that the major categories of hits from the bright screen, factors required for mitochondrial function and translation, are unified by a common mechanism: reducing ribosome collisions at inefficiently decoded selenocysteine codons.

### EEF2K does not mediate selenoprotein induction during oxidative stress

In the presence of stressors such as nutrient starvation or hypoxia, the key translation elongation factor EEF2 is known to be phosphorylated at threonine 56 by EEF2 kinase (EEF2K), resulting in its inhibition (*62*). We therefore tested whether EEF2K was responsible for the slowing of elongation and the consequent increase in selenoprotein production during oxidative stress. Phosphorylation of EEF2 was not increased upon pharmacologic inhibition of complex I, III or V (**Fig. 5E**). Moreover, treatment of GPX1 and GPX4 reporter cells with the EEF2K inhibitor A-484954 (EEF2Ki), which effectively blocked EEF2 phosphorylation (**Fig. 5F**), had no effect on baseline reporter expression nor reporter induction upon ETC inhibition with oligomycin A (OMA) (**Fig. 5, G and H**). To definitively rule out a role for EEF2K in regulation of selenoprotein production during oxidative stress, clonal *EEF2K* knockout cell lines were generated which lacked any detectable EEF2 phosphorylation (**Fig. 5I**). GPX1 and GPX4 protein levels increased equivalently in wild type and *EEF2K* knockout cells upon loss of ETC components or TXNRD1 (**Fig. 5, I to K**), excluding the possibility that this response is mediated by EEF2K.

### Oxidized glutathione binds EEF1G and triggers disassembly of the EEF1H complex

Our finding that EEF2K does not mediate the upregulation of selenoprotein synthesis during oxidative stress prompted us to examine other components of the translation elongation machinery for potential sensors of the cellular redox state. Interestingly, multiple components of the EEF1H complex, which recycles EEF1A-GDP to its GTP bound form after tRNA delivery, contain partial or complete glutathione S-transferase (GST) domains (**Fig. 6A**). EEF1H is a high molecular weight assembly comprising EEF1A, the guanine nucleotide exchange factors (GEFs) EEF1B and EEF1D, a structural subunit EEF1G, as well as valine tRNA synthase (VARS) (*63–65*). While EEF1B and VARS contain only GST-C-terminal (GST-C) domains that mediate substrate binding in canonical GSTs, EEF1G harbors an additional GST-N-terminal (GST-N) domain that directly binds glutathione (*66, 67*). These observations raised the possibility that glutathione might directly regulate the function of EEF1G. We note that ETC inhibition and TXNRD1 loss both impact the ratio of reduced to oxidized glutathione (GSH:GSSG; **fig. S7F**), highlighting the potential of the glutathione system to act as a common readout of these distinct perturbations which both increased selenoprotein production.

**Figure 6.**
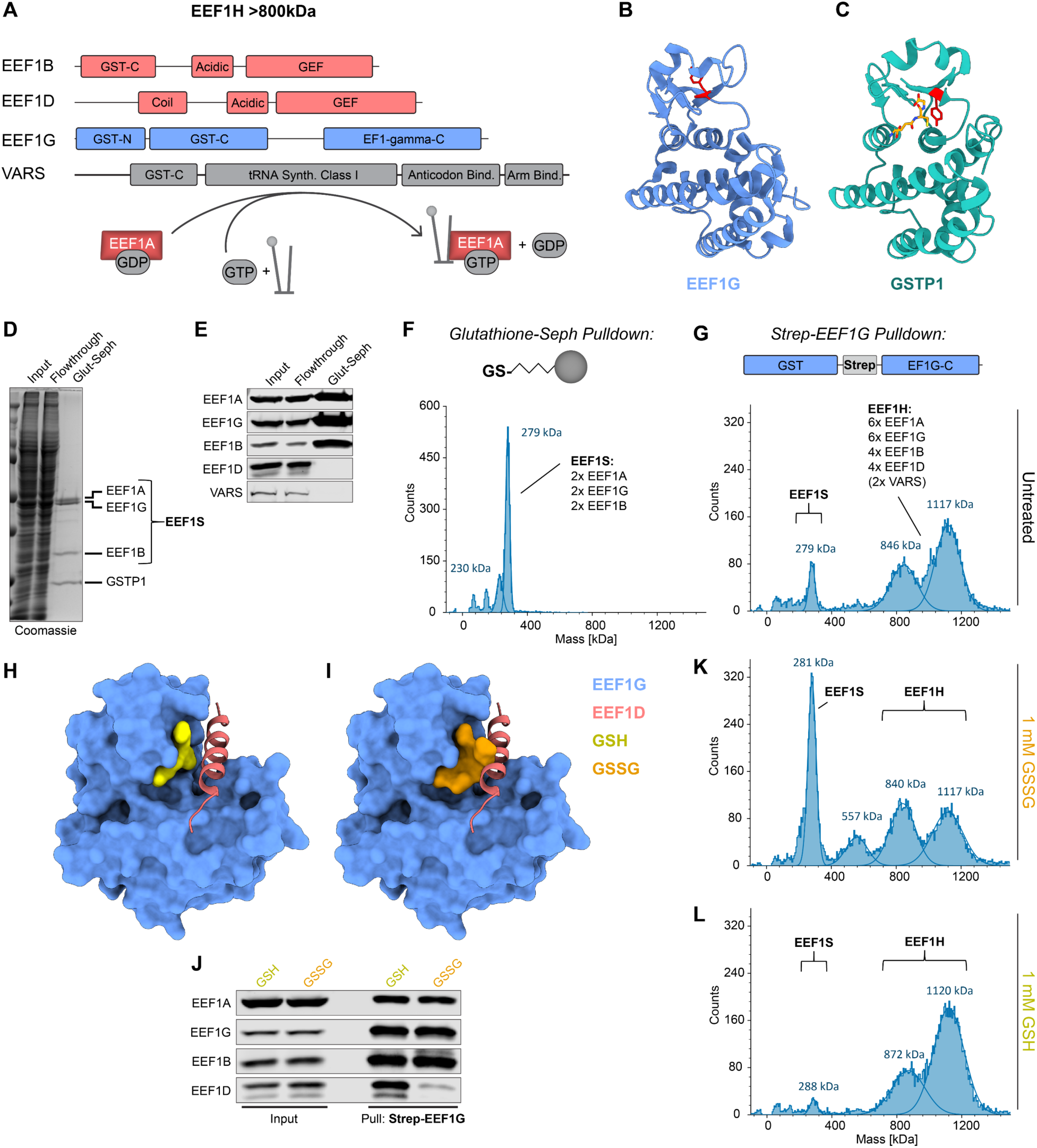
Binding of oxidized glutathione to EEF1G leads to EEF1H disassembly. (**A**) Domain organization of the subunits of the EEF1H complex, which recycles EEF1A-GDP to EEF1A-GTP, enabling tRNA delivery during translation. (**B**) and (**C**) Structures of the GST domain of human EEF1G (PDB: 5JPO) (B) and GSTP1 bound to reduced glutathione (PDB:19GS) (C). Red, tyrosine 7 in EEF1G and catalytic tyrosine 8 in GSTP1; yellow, glutathione. (**D**) to (**F**) Glutathione-Sepharose pulldown from HEK293T lysates. Recovered proteins were analyzed by Coomassie staining (D), western blotting (E), or mass photometry (F). The predicted stoichiometry of the EEF1S complex, based on molecular weight of components and Coomassie staining, is indicated on the mass photometry plot. (**G**) Mass photometry of the EEF1H complex, recovered by affinity purification of endogenous strep-tagged EEF1G from HEK293T lysates. The predicted stoichiometry of the EEF1H complex, based on molecular weight of components and Coomassie staining, is indicated. (**H**) and (**I**) Structure of the EEF1G GST domain (blue; aa 1-226) in complex with the N-terminal helix of EEF1D (red; aa 11-76) (PDB: 5JPO) superimposed with AF3 modeled binding of reduced (yellow) (H) or oxidized (orange) (I) glutathione. (**J**) Western blot analysis of strep-EEF1G pulldown from HEK293T cells in the presence of 1 mM reduced (GSH) or oxidized (GSSG) glutathione. (**K**) and (**L**) Mass photometry of purified EEF1H incubated with either 1 mM oxidized (K) or reduced (L) glutathione before measurement.

The structure of the EEF1G GST domain shows a similar overall topology compared to the GST-N and GST-C domains of canonical GSTs (**Fig. 6, B and C**) (*68*). However, the inferred catalytic residue for glutathione transfer in EEF1G, Y7, is facing away from the glutathione binding site, and previous studies reported weak or undetectable GST activity of EEF1G orthologs (*67, 69–71*). Thus, glutathione binding, not conjugation, likely represents the main function of the EEF1G GST-N domain. To examine the composition of glutathione-bound EEF1G complexes, we used glutathione-Sepharose to recover glutathione-binding proteins from HEK293T cell lysates. These pulldowns retrieved the most abundant canonical GST in this cell type, GSTP1, as well as an EEF1 subcomplex consisting of EEF1G, EEF1A, and EEF1B (**Fig. 6, D and E**). We further characterized this smaller glutathione-associated EEF1 complex, which we termed EEF1S, by size exclusion chromatography (SEC) and determined its molecular weight by mass photometry (**Fig. 6F and fig. S9, A and B**). Based on its molecular weight of approximately 270 kDa and the 1:1:1 relative stoichiometry of each subunit estimated from Coomassie staining, this EEF1S complex most likely represents a dimer of EEF1A, EEF1G, and EEF1B. Canonical EEF1H, which additionally contains EEF1D and VARS, was prepared by affinity purification of endogenously Strep-tagged EEF1G from HEK293T cells (**fig. S9, C and D**). Mass photometry of this complex, coupled with estimation of its stoichiometry by Coomassie staining, revealed a likely subunit stoichiometry of six copies of EEF1G and EEF1A, four copies of EEF1B and EEF1D, with and without two copies of VARS (∼1100 kDa and ∼820 kDa respectively; **Fig. 6G**).

To gain insight into the assembly of EEF1S and EEF1H, and the potential regulation of these complexes by glutathione, we examined a previously determined structure of the EEF1G GST domain in complex with an N-terminal fragment of EEF1D (**Fig. 6H**). In this structure, an alpha helix of EEF1D (residues 14-28) binds to EEF1G in close proximity to the glutathione binding site. Mutagenesis of key residues that are predicted to be important for this interaction in either EEF1D (E22 and Y26) or EEF1G (R14) strongly diminished the interaction, confirming this EEF1G:EEF1D binding interface (**fig. S9, E to G**). We next used AlphaFold3 (AF3) (*72*) to model how glutathione binding impacts this interaction. Interestingly, while the binding of reduced glutathione (GSH) to EEF1G was compatible with the EEF1D interaction in the AF3 model (**Fig. 6H**), oxidized glutathione (GSSG) was predicted to sterically clash with EEF1D in this binding pocket (**Fig. 6I**). Indeed, pulldown of Strep-tagged EEF1G in the presence of oxidized glutathione greatly decreased co-purification of EEF1D compared to pulldowns performed in the presence of reduced glutathione (**Fig. 6J**). Furthermore, when oxidized glutathione was added to EEF1H, we observed that the complex disassembled into a smaller complex that lacked EEF1D and was indistinguishable from EEF1S in mass (∼270kDa; **Fig. 6K and fig. S9H and I**). Addition of reduced glutathione to purified EEF1H had the opposite effect, decreasing the basal amount of EEF1S detectable by mass photometry (**Fig. 6L**). Notably, the exclusive recovery of EEF1S by glutathione-Sepharose pulldown suggested that the aliphatic linker in the glutathione-resin likely mimicked binding of oxidized glutathione and sterically blocked EEF1D binding. These data therefore documented that the binding of oxidized glutathione to EEF1G induces dramatic reorganization of EEF1H and loss of a core subunit of the complex.

### Glutathione binding to EEF1G is required for slowed translation and selenoprotein induction during oxidative stress

Our finding that oxidized glutathione, which accumulates during oxidative stress, impacts the organization and composition of the EEF1H complex suggested a mechanism that may couple the efficiency of translation elongation to the cellular redox state. To investigate this possibility, we aimed to identify an EEF1G mutant that lost the ability to bind glutathione but maintained canonical EEF1G function. Analysis of EEF1G-GST-N domain conservation among eukaryotes and comparison of the domain to canonical human GSTs identified EEF1G^P59^ as the only amino acid that was fully conserved among EEF1G orthologs and present in all human GSTs (**fig. S10. A and B**). EEF1G^P59^ exists in a cis-conformation that organizes the structure of an extended loop in the GST-N domain that is necessary for glutathione binding (**Fig. 7A**) and has been reported to be essential for catalytic activity of canonical GSTs (*73*).

**Figure 7.**
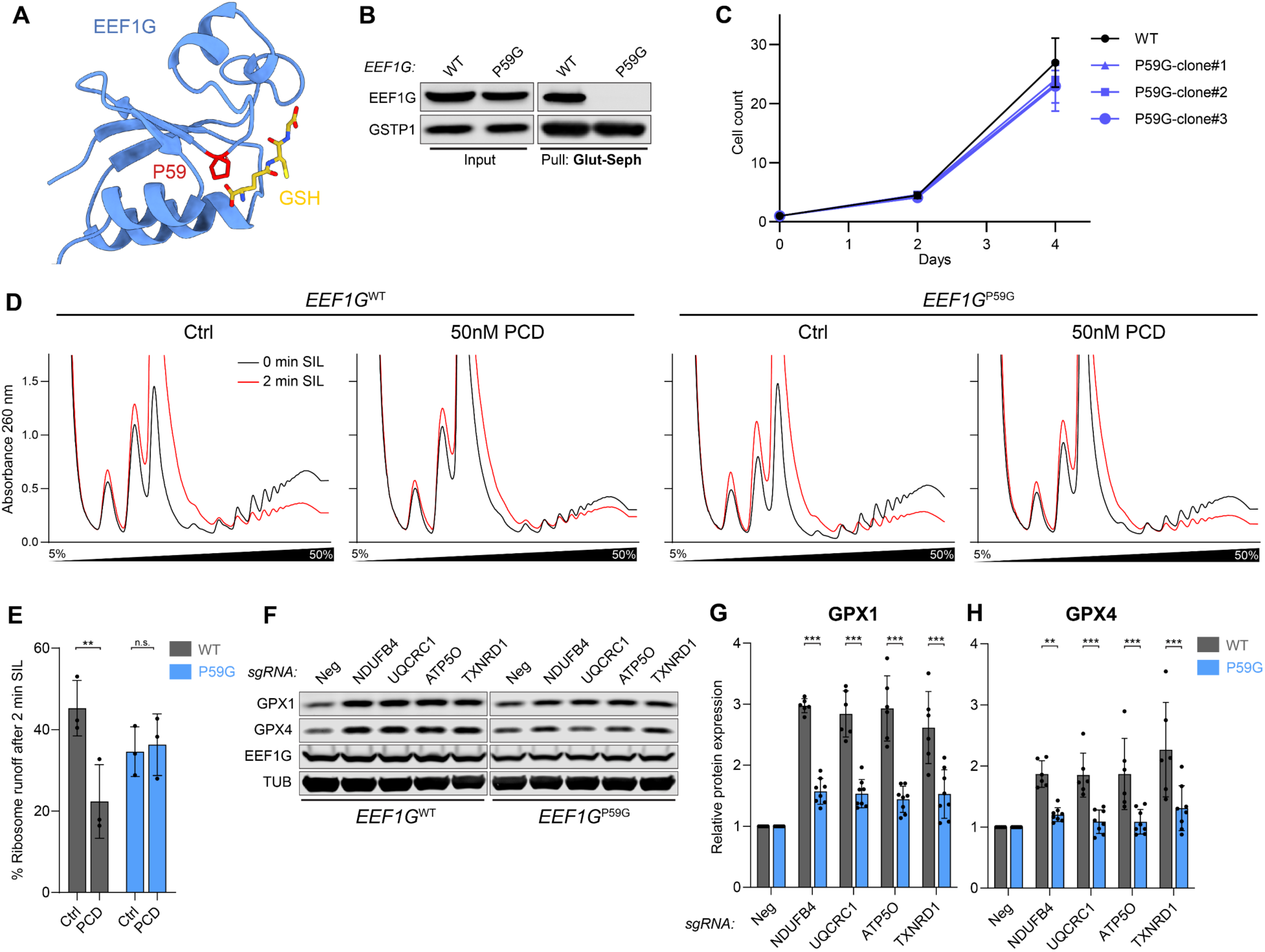
Glutathione binding by EEF1G is required for slowed elongation and increased selenoprotein synthesis during oxidative stress. (**A**) AF3 model of EEF1G in complex with GSH showing conserved cis-proline at position 59 (red). (**B**) Western blot analysis of glutathione-Sepharose pulldowns from *EEF1G*^WT^ and *EEF1G*^P59G^ HCT116 cells. (**C**) Proliferation of parental HCT116 cells (WT) and three independent *EEF1G*^P59G^ homozygous knock-in clones. (**D**) and (**E**) Ribosome runoff experiments (D) and associated quantification (E) in *EEF1G*^WT^ or *EEF1G*^P59G^ HCT116 cells in in the presence or absence of PCD. Polysome abundance was monitored by sucrose gradient sedimentation before and after 2 minutes of treatment with the translation initiation inhibitor silvestrol (SIL; 2 µM). Area under the curve for gradient positions with polysomes (fractions 8-12) was quantified and normalized to 0 min SIL condition. n=3 biological replicates. *P* values were calculated by 2-way ANOVA with Šídák’s multiple comparisons post-test. n.s., not significant; **p < 0.01. (**F**) to (**H**) Western blot analysis (F) and associated quantification (G and H) of GPX1 and GPX4 protein levels in *EEF1G*^WT^ or *EEF1G*^P59G^ HCT116 cells transduced with lentiviral constructs encoding Cas9 and the indicated sgRNAs. Protein abundance was normalized to EEF1G and to sgNeg. Mean ± SD plotted. n=6 biological replicates for *EEF1G*^WT^ cells and n=8 biological replicates for *EEF1G*^P59G^ cells from 3 independent knock-in clones. *P* values were calculated by 2-way ANOVA with Tukey’s multiple comparisons post-test. **p < 0.01, ***p < 0.001.

CRISPR-mediated genome editing was used to generate clonal HCT116 cell lines with homozygous P59G mutations at the endogenous *EEF1G* locus. The P59G mutation had no effect on EEF1G protein abundance and mutant cell lines grew normally (**Fig. 7, B and C**). However, glutathione binding of EEF1G^P59G^ was fully abolished (**Fig. 7B**). We then performed polysome run-off assays to assess translation elongation rates in wild type or *EEF1G*^P59G^ cells in the presence of oxidative stress. In contrast to wild type cells, which exhibited greatly decreased run-off following treatment with the ROS-inducing compound PCD, translation elongation was relatively preserved in *EEF1G*^P59G^ cells (**Fig. 7, D and E**). Moreover, enhanced production of selenoproteins GPX1 and GPX4 was abrogated in *EEF1G*^P59G^ cells following loss of ETC complex components or TXNRD1 (**Fig. 7, F to H**). Altogether, these data support a model in which EEF1G, through its ability to bind glutathione, couples the cellular redox state to the rate of translation elongation, thereby enhancing production of detoxifying selenoproteins under conditions of oxidative stress.

## Discussion

Although the core factors required for selenocysteine decoding were discovered more than two decades ago (*15–17, 19, 20, 22, 25*), a systematic investigation of this pathway using unbiased genetic approaches has not previously been reported. This is likely due to the absence of the selenocysteine pathway in yeast, a model organism of choice for genetic screens. Here, we leveraged a CRISPR-based screening approach to interrogate the selenocysteine pathway directly in human cells, enabling the comprehensive identification of regulators of this process. From these studies, we uncovered two mechanisms that govern selenocysteine incorporation and couple the production of detoxifying selenoproteins to the redox state of the cell. First, we found that selenoprotein production is limited by ribosome collisions that occur at slowly decoded selenocysteine codons. Consequently, any perturbation that slows translation, such as depletion of core translation elongation factors, results in an enhanced window for selenocysteine decoding, which mitigates collisions and augments selenoprotein production.

Second, we documented that oxidative stress slows elongation in mammalian cells, thereby increasing selenoprotein synthesis through this mechanism. Together, these events form the basis of a homeostatic feedback mechanism that enables a rapid increase in selenoprotein translation during oxidative stress by reducing ribosome collisions on these transcripts.

Through these studies, the core translation elongation factor EEF1G emerged as a key sensor of the cellular redox state that slows elongation during periods of oxidative stress. EEF1G has primarily been viewed as a structural subunit of the EEF1H complex, which exchanges GTP for GDP on the tRNA delivery factor EEF1A. Over 30 years ago, EEF1G was noted to possess a glutathione S-transferase (GST)-like domain (*66, 70*), but in the absence of robust GST catalytic activity (*67, 70, 71*), the role of glutathione binding by EEF1G has remained unclear. Our findings have provided an answer to this longstanding question by demonstrating that EEF1G acts as a sensor for oxidized glutathione (GSSG), a metabolite whose levels specifically increase during oxidative stress (*74*) or upon deficiency of cellular reducing potential (*75*). For a sensor of oxidative stress, detection of oxidized glutathione is advantageous since reduced glutathione (GSH) is highly abundant, and its substantial depletion is only expected to occur under extreme conditions. We documented that the binding of oxidized glutathione to EEF1G induces a dramatic reorganization of the EEF1H complex into a lower molecular weight form that lacks the EEF1D subunit, and which presumably recycles GTP-bound EEF1A less efficiently. While additional biochemical and structural studies are necessary to determine how the organization of EEF1H, and its subcomplex EEF1S, impacts the activity of these assemblies, our data reveal that the sensing of oxidized glutathione by EEF1G can effectively couple the redox state of the cell to the translation elongation rate.

These findings add a new dimension to the growing link between the translation machinery and cellular stress responses. During oxidative stress, ribosomes may stall stochastically due to damage to mRNAs, tRNAs, or ribosomes, leading to widespread collisions (*2, 3*). The slowing of translation elongation by EEF1G under these conditions would be expected to limit these collisions, providing a window for ribosome quality control (RQC) pathways to remove stalled ribosomes while cellular stress is resolved and damaged mRNAs and translation components are cleared. Consequently, full activation of the RSR, and downstream cell death pathways, may be avoided when stress is transient. Importantly, experimental evidence that slowed elongation limits RSR activation was provided by a genome-wide CRISPR screen which showed that loss of genes required for canonical A-site decoding, including hits recovered in our screen such as *EEF1G* and *EEF1A*, enhanced cell survival in the setting of global collisions induced by the elongation inhibitor ANS (*7*). Thus, sensing of oxidized glutathione levels by EEF1G may similarly raise the threshold for full RSR activation, providing a first line of defense against redox perturbations. Oxidative stress can further activate the ISR (*1*), leading to a global reduction in translation initiation. By reducing ribosome load across the transcriptome, this effect is expected to synergize with EEF1G-mediated regulation of elongation rate to further limit collisions and restrict full activation of the RSR to settings where activation of cell death is inescapable.

In addition to globally reducing translation initiation, the ISR selectively enhances the translation of specific factors that contribute to the management of broad perturbations such as protein misfolding, nutrient deprivation, and oxidative stress (*1*). A key feature that enables ISR-mediated upregulation is the presence of one or more upstream open reading frames (uORFs) in the 5′ UTRs of transcripts that encode ISR-induced proteins (*76–78*). Analogously, our findings suggest that programmed ribosome collisions confer the selective regulation of key transcripts by pathways that modulate translation elongation. This concept is exemplified by selenoprotein synthesis, which is limited by ribosome collisions that occur at slowly-decoded selenocysteine codons. The rheostatic control of translation elongation rate through EEF1G-mediated redox sensing is coupled to selenoprotein production through these collisions, ultimately allowing an immediate boost in selenoprotein levels during oxidative stress. Beyond selenoprotein-encoding transcripts, ribosome collisions occur at defined sites throughout the transcriptome (*45, 79–81*), pointing to broader control of translation by EEF1G-mediated redox sensing. Features that induce collisions include sequences that encode amino acids that are poor substrates for peptide bond formation, such as proline and glycine, or stretches of positively-charged amino acids such as lysine, which can slow translation by interacting with the negatively-charged ribosome exit channel (*45, 79–82*). Intriguingly, global disome profiling revealed that many transcripts encoding factors that participate in oxidation-reduction reactions, including numerous proteins that do not contain selenocysteine, exhibit strong collision signatures (*45*). Thus, the slowing of translation elongation by EEF1G during oxidative stress may broadly reprogram the translatome to promote redox balance.

The selective regulation of specific sets of transcripts by programmed ribosome collisions is likely not limited to oxidative stress conditions. For example, we observed that transcripts encoding core translation elongation factors, such as EEF1A and EEF1G, exhibit strong disome and ZAKα peaks (*7, 45*) (**fig. S11, A and B**), suggesting that the synthesis of these factors is limited by ribosome collisions. This scenario provides a mechanism for rapid enhancement of elongation factor production when translation elongation slows, thereby allowing precise control of the levels of these factors to meet cellular demand. Given the frequency of ribosome collisions detected throughout the transcriptome (*7, 45, 79–81*), it is likely that many more proteins are regulated in this manner. Thus, the homeostatic coupling of protein production to translation elongation rate via colliding ribosomes may be a widespread mechanism for gene regulation.

## Materials and methods

### Cell culture

Cell lines were obtained from the American Type Culture Collection (ATCC) and verified to be free of mycoplasma contamination. HCT116 and HEK293T cells were cultured in Dulbecco’s Modified Eagle’s Medium (DMEM) with high glucose and pyruvate (Corning) supplemented with 10% (v/v) fetal bovine serum (FBS; Sigma) and 1x Penicillin and Streptomycin (Sigma). During transfection and lentiviral transduction, cells were cultured in antibiotic-free medium.

### *GPX*-eGFP reporter cell line generation

Gene loci of *GPX1* and *GPX4*, spanning the 5′ UTR to the end of the coding sequence (CDS), were amplified from human genomic DNA (gDNA) using Primestar GLX proofreading polymerase (Takara). These amplicons were cloned together with a synthetic gene fragment encoding enhanced green fluorescent protein (eGFP) and the *GPX1* or *GPX4* 3′ UTR (IDT) using NEBuilder HiFi DNA Assembly (New England Biolabs) into a modified PiggyBac (PB) donor backbone (XLone) (*34*) under control of the EF1α promoter. The GPX4 reporter plasmid containing a self-cleavable peptide sequence was generated using the same strategy, incorporating a synthetic fragment with a PT2A sequence between *GPX4* and eGFP. The GPX4M reporter construct was created by site-directed mutagenesis of the selenocysteine codon (TGA) to a cysteine codon (TGT) in the GPX4 reporter plasmid. Primer and synthetic gene fragment sequences are provided in **table S3**. For stable reporter cell line generation, 600 ng of PB donor construct DNA and 200 ng of pCAGEN-HyPBase transposase vector were reverse-transfected into HCT116 cells in each well of a 6-well plate using 2 µL FuGene HD (Promega) following the manufacturer’s protocol. After 12 days of culture to ensure genomic integration, GFP-positive single cells were sorted into 96-well plates, and clonal cell lines were established.

### Flow cytometry

Cells were washed with phosphate-buffered saline (PBS), incubated with 0.05% Trypsin-EDTA (Thermo Scientific) at 37°C for 3 minutes, and collected in DMEM supplemented with 10% FBS. After centrifugation at 200 x g for 3 minutes, the supernatant was removed and cell pellets were resuspended in flow buffer (3% FBS, 2 mM EDTA in PBS). Samples were analyzed using an Accuri C6 flow cytometer (BD Biosciences), collecting data from at least 20,000 live cells per sample, as determined by forward and side scatter. Flow cytometry data were processed and visualized using FlowJo software (version 10.10.0; BD Biosciences).

### Lentiviral CRISPR-Cas9-mediated gene knockout

Single guide RNA (sgRNA) sequences targeting genes of interest were cloned into the lentiCRISPRv2 backbone (Addgene #52961) using NEBuilder HiFi DNA Assembly (New England Biolabs). For lentivirus production, 1.2 x 10^6^ HEK293T cells were seeded per well in 6-well plates. The following day, cells were transfected with 1 μg lentiCRISPRv2, 0.6 μg psPAX2 (Addgene #12260), and 0.4 μg pMD2.G (Addgene #12259) using 5 μL FugeneHD transfection reagent (Promega). After 24 hours, the medium was replaced, and viral supernatant was collected for another 24 hours, filtered through a 0.45 μm membrane, and stored in aliquots at −80°C. Target cells were transduced with lentiviral supernatant in the presence of 5 μg/ml polybrene. Two days post-transduction, cells were transferred to medium containing 1 μg/ml puromycin for selection. Knockout cell pools were maintained under puromycin selection for 5–7 days prior to downstream analyses.

### CRISPR-Cas9 genome-wide knockout screen for selenocysteine decoding

Genome-wide CRISPR-Cas9 knockout screens were performed as previously described (*83*). Two independent clones for each reporter were used. A total of 1.4 x 10^8^ cells were infected with the Brunello pooled lentiviral CRISPR library (Addgene #73179) in the presence of 5 μg/ml polybrene at a multiplicity of infection (MOI) between 0.3 to 0.5. Two days post-transduction, selection with 1 μg/ml puromycin was initiated. Cells were passaged every 48-72 hours, maintaining a minimum of 5 x 10^7^ cells per passage to maintain >500X coverage of the library.

After 8 days, puromycin concentration was reduced to 0.5 μg/ml. After 10 days of puromycin selection, the brightest and dimmest 0.75% of eGFP expressing cells from each reporter clone were collected using a FACS Melody cell sorter (BD Biosciences). A starting pool of at least 4 x 10^7^ live cells were sorted per clone to maintain >500X coverage of the library, yielding at least 3 x 10^5^ cells in each collected gate. gDNA from sorted cells was extracted by phenol-chloroform extraction. gDNA from 6 x 10^7^ unsorted cells was isolated with the Masterpure DNA isolation kit (Lucigen) following the manufacturer’s instructions. Sequencing libraries were prepared from the isolated gDNA using two rounds of PCR amplification with Herculase II DNA polymerase (Agilent). For unsorted cells, 4 μg of gDNA was used in each 100 μl reaction for PCR 1, with 57 reactions used in total for each sample. For sorted cells, all recovered gDNA was amplified in a single 100 μl reaction. After 18 cycles of initial PCR amplification, unsorted reactions were pooled and 5% of reaction products were used for a second PCR reaction with 10 cycles to introduce Illumina sequencing adapters and barcodes. Primer sequences are provided in **table S3**. Final PCR products were purified using AMPure XP beads (Beckman Coulter). Libraries were pooled and sequenced on a NextSeq 2000 (Illumina) instrument with 100 bp single end reads at an average depth of 2.2 x 10^7^ reads per library. sgRNA sequences were extracted from fastq files through an in-house Galaxy script (*84*) and normalized reads counts were calculated. Gene-level enrichment in sorted versus unsorted populations was determined using MAGeCK (*85*).

### Western blotting

Cells were washed once with ice-cold PBS and lysed in polysome buffer (1% Triton X-100, 150 mM KCl, 20 mM Tris-HCl pH 7.4, 15 mM MgCl_2_, 1X Calbiochem EDTA free proteinase inhibitor cocktail III), briefly vortexed, and kept on ice for 10 minutes. When protein phosphorylation was analyzed, PhoSTOP (Roche) was added to the lysis buffer. Lysates were centrifuged at 18,000 g for 5 minutes at 4°C and supernatant was stored at-80°C until use. Proteins were separated by electrophoresis on 10% to 12% SDS Tris-glycine polyacrylamide gels and wet-blotted onto 0.45 μm nitrocellulose membranes. Membranes were blocked in PBS with 0.1% Tween-20 (PBST) containing 5% non-fat milk for 1 hour at room temperature, briefly washed in PBST, and incubated with primary antibodies in PBST containing 3% BSA overnight at 4°C. Blots were then washed three times in PBST and incubated with fluorescent secondary antibodies (LI-COR) in PBST with 5% non-fat milk for one hour at room temperature. After three washes with PBST, images were acquired on an Odyssey fluorescent western blot imaging system (LI-COR Biosystems). Antibodies used for western blot are listed in **table S4**.

### Ribosome fractionation

Two days prior to the experiment, HCT116 cells were seeded to reach approximately 75% confluency on the day of fractionation. One 15 cm plate was seeded per sample. One day before the experiment 5-50% continuous sucrose gradients (prepared in 150 mM KCl, 20 mM Tris-HCl pH 7.4, 5 mM MgCl_2_) were prepared using a BioComp Gradient Master and stored at 4°C. On the day of the experiment, cells were treated with translation elongation inhibitors when appropriate (cycloheximide, anisomycin, didenmin B, or ternatin), and then immediately placed on ice, washed once with ice-cold PBS (containing the respective translation elongation inhibitor when appropriate), and collected in PBS with a rubber cell scraper. Cells were pelleted by centrifugation at 500 g for 2 minutes at 4°C. Pellets were resuspended in 450 μl polysome lysis buffer (1% Triton X-100, 150 mM KCl, 20 mM Tris-HCl pH 7.4, 15 mM MgCl_2_) supplemented with EDTA-free proteinase inhibitor cocktail III (Calbiochem), 200 u/ml recombinant RNasin (Promega), and the respective translation elongation inhibitor when appropriate. Lysates were centrifuged at 18,000 g for 5 minutes at 4°C, the supernatant was transferred to a new tube, and 40 μl was removed as input control. 400 μl of the remaining sample was then gently layered onto a prepared sucrose gradient and centrifuged at 38,000 rpm in a TH-641 rotor for 2 hours at 4°C using a Sorvall WX (Thermo Scientfic) ultracentrifuge with acceleration and deceleration settings of 8 and 0, respectively. After centrifugation, gradients were loaded into a piston-based BioComp ribosome fractionator. Absorbance at 260 nm was recorded at a piston speed of 0.2 mm/s, and 12 fractions were collected per gradient. Proteins were precipitated from fractions by adding trichloroacetic acid (TCA) to a final concentration of 12.5% and incubation on ice for 30 minutes. Protein precipitates were pelleted by centrifugation at 12,000 g for 10 minutes at 4°C. Protein pellets were washed twice with acetone and resuspended in 2X Laemmli buffer (2% SDS, 10% glycerol, 0.02% bromophenol blue, 0.05 M Tris-HCl pH 6.8, 5% 2-mercaptoethanol) for analysis by SDS-PAGE and western blot.

### Quantification of cellular ROS

HCT116 cells were transduced with lentiCRISPRv2 lentiviral constructs expressing Cas9 and sgRNAs. Following 6 days of puromycin selection (1 μg/ml), cells were stained with the ROS-ID Total ROS detection kit (Enzo Life Sciences) according to the manufacturer’s instructions. As a positive control, untransduced HCT116 cells were treated for 30 minutes with 100 μM pyocyanin, a ROS-inducing agent. Stained cells were analyzed using an Accuri C6 flow cytometer as described in the flow cytometry section.

### Quantification of cellular GSH/GSSG ratio

Following transduction with lentiCRISPRv2 constructs and four days of puromycin selection (1 μg/ml), HCT116 cells were seeded into non-transparent, tissue culture-treated flat-bottom 96-well plates and cultured for an additional two days. For each condition, two wells were used for total glutathione quantification (GSH+GSSG) and two wells for oxidized glutathione (GSSG) measurement. Cells were lysed in-well in 1X Passive Lysis buffer (Promega) and the ratio of reduced to oxidized glutathione was determined using the GSH/GSSG-Glo Assay (Promega) according to the manufacturer’s instructions.

### RNA isolation and qRT-PCR

Total RNA was isolated with the RNeasy kit (Qiagen) including the optional on-column DNase digestion step according to the manufacturer’s instructions. 1 µg of RNA was reverse transcribed using the PrimeScript RT MasterMix (Takara) in a final volume of 20 μl. The resulting cDNA was diluted 1:3 with RNase free water and 1 μl was used in a 10 μl qRT-PCR reaction with Power SYBR Green (Thermo Scientific) on a Quant Studio 5 qPCR instrument (Thermo Scientific). Transcript levels were quantified using the ΔΔCt method and primer amplification efficiencies were validated using serial dilutions. Primer sequences are provided in **table S3**.

### Polysome runoff assays

Cells and gradients were prepared as described in the ribosome fractionation section. 1 μM silvestrol (MedChemExpress) was added to cells growing in 15 cm dishes. Plates were gently rocked to evenly distribute the compound and immediately placed back into the 37°C incubator for 2 minutes. Cycloheximide was then added to non-runoff control and silvestrol-treated plates at a final concentration of 100 μg/ml to halt translation elongation. Plates were quickly rocked to disperse cycloheximide, placed on ice, and processed immediately for ribosome fractionation. PBS used for washing and polysome lysis buffer were both supplemented with 100 μg/ml cycloheximide.

### Generation of clonal knockout cell lines using CRISPR-Cas9

HCT116 or HEK293T cells were reverse transfected with 800 ng of pX458 plasmid expressing eGFP, Cas9, and the desired sgRNA (see **table S3** for oligonucleotide sequences) in 6-well plates using FuGene HD transfection reagent (Promega). Two days post-transfection, cells were passaged into new tissue culture plates and four days post-transfection, eGFP-positive single-cells were sorted into 96 well plates using a BD FACS Melody cell sorter. Clonal cell lines were established from individual wells and gene knockout was confirmed by western blot.

### Generation of knock-in cell lines using CRISPR-Cas9

Endogenous knock-in of tags or point mutations was carried out by nucleofection of CRISPR mRNP containing Cas9 nuclease, sgRNA, and single stranded donor oligonucleotide for homology directed repair (HDR). All reagents for gene editing were obtained from IDT (Alt-R-sgRNA, Alt-R-HDR donor oligo, AltR-Spy-Cas9 nuclease V3, Alt-R-HDR enhancer V2, Alt-R-electroporation enhancer). HDR donors were 200 nucleotides in length and were designed with left and right homology arms of approximately similar length. In cases where HDR editing did not disrupt the sgRNA binding site, the PAM sequence in the HDR donor was mutated to prevent Cas9 recutting of edited alleles. Sequences of HDR donors and sgRNAs are provided in **table S3**. Two days prior to editing, cells were seeded so they reached approximately 70% confluency on the day of nucleofection. Cas9-sgRNA mRNP was assembled by mixing 2 μl PBS, 7 μl of sgRNA (50 μM), and 6 μl Cas9 nuclease (61 μM) followed by incubation for 15 minutes at room temperature. In parallel, HDR donor mixes were prepared by combining 4.2 μl PBS, 7.2 μl donor oligo (50 μM), and 3.6 μl electroporation enhancer (50 μM). Cells from one 10 cm dish were trypsinized, collected in DMEM media containing 10% FBS, centrifuged at 200 g for 3 minutes, and resuspended in 1 ml PBS. After counting, 1.3 x 10^6^ cells were pelleted again and gently resuspended in 70 μl Lonza electroporation solution (HEK293T: solution SF; HCT116: solution SE) supplemented with 80 ng of pMax-eGFP vector (Addgene #177825). Cell suspensions were gently mixed with mRNP and HDR mixtures and transferred to a Lonza nucleofection cuvette. Electroporation was performed using a Lonza 4D Nucleofector with program EN-113 (HCT116) or DS-150 (HEK293T). Following nucleofection, 400 μl of 37°C pre-warmed DMEM containing 10% FBS was added to the cuvette and electroporated cells were gently transferred out of the cuvette to a 6 well plate containing 2 ml of prewarmed medium and 3 μl of HDR enhancer. Medium was replaced the following day, and cells were passaged two days post-nucleofection. 4 to 7 days after nucleofection, single eGFP-positive cells were sorted into 96 well plates using a BD FACS Melody cell sorter and clonal cell lines were established.

### Generation of lentiviral *EEF1G* and *EEF1D* expression constructs

The coding sequence (CDS) of *EEF1G* was PCR-amplified from cDNA generated from parental HEK293T cells (for untagged *EEF1G*) or from CRISPR-Cas9-edited HEK293T cells containing an internal Twin-Strep tag inserted in the unstructured region between the GST and C-terminal domains. PCR fragments were cloned using NEBuilder HiFi DNA Assembly (New England Biolabs) into a minimal lentiviral backbone derived from Lenti-Cas9-Blast (Addgene #52962), in which the Cas9 and blasticidin resistance cassettes had been removed by digestion with XbaI and EcoRI. *EEF1G* R14E was generated by digestion of Lenti-EEF1G with AgeI and XbaI and HiFi reaction with a synthetic gene fragment carrying the mutation and a PCR product encoding the remaining part of *EEF1G*.

*EEF1D* CDS was amplified by PCR from HCT116 cDNA and cloned with NEBuilder HiFi DNA Assembly (New England Biolabs) into a modified pLV-EF1a-IRES-Blast (Addgene # 85133) backbone, in which a 3XFLAG tag was inserted downstream of the EF1α promoter. *EEF1D* E22R and Y26E were generated by PCR-based site-directed mutagenesis. Sequences of oligonucleotides and synthetic gene fragments are provided in **table S3**.

### Affinity purification and size exclusion chromatography of EEF1 complexes

Approximately 5 x 10^8^ HEK293T cells were washed twice with ice-cold PBS and resuspended in 8 ml Co-IP lysis buffer (0.5% Triton X-100, 25 mM HEPES pH 7.4, 150 mM KOAc, 5 mM NaCl, 2 mM Mg(OAc)_2_, 15% glycerol, 1X EDTA-free protein inhibitor cocktail III). Cells were lysed by passing the suspension twice through a 26G needle. Lysates were centrifuged at 14,000 x g for 10 minutes at 4°C, and the supernatant was transferred to a new tube. Input sample was removed, and affinity pulldowns were performed by incubating the lysate with 250 μl of washed StreptactinXT or Glutathione Sepharose 4B resin (Cytiva) for 2 hours at 4°C with rotation.

Following incubation, beads were washed with low-detergent wash buffer (0.025% Triton X-100, 25 mM HEPES pH 7.4, 150 mM KOAc, 5 mM NaCl, 2 mM Mg(OAc)_2_, 5% glycerol), resuspended in wash buffer, and applied to a gravity-flow column. Columns were then washed with no-detergent wash buffer (25 mM HEPES pH 7.4, 150 mM KOAc, 5 mM NaCl, 2 mM Mg(OAc)_2_, 5% glycerol) and proteins were eluted by two sequential additions of 4 ml elution buffer (25 mM HEPES pH 7.4, 25 mM Tris pH 8, 150 mM KOAc, 5 mM NaCl, 2 mM Mg(OAc)_2_) containing either 10 mM reduced glutathione for the glutathione Sepharose pulldown or 20 mM biotin for the StreptactinXT pulldown. Eluates were concentrated to approximately 550 μl using Pierce PES protein concentrator spin columns (Thermo Scientific) with a 30 kDa molecular weight cutoff. For size exclusion chromatography (SEC), 500 μl of concentrated eluate was injected into an ÄKTA Pure Fast Protein Liquid Chromatography (FPLC) system (Cytiva) and separated on Superdex 200 SEC columns equilibrated with SEC buffer (25 mM HEPES pH 7.4, 25 mM Tris pH 8.0, 150 mM KOAc, 5 mM NaCl, 2 mM Mg(OAc)₂). Elution fractions were collected at a fixed volume of 0.5 ml.

### 3X-FLAG immunoprecipitation

For each sample, 75 μl of Protein-G-coupled Dynabeads (Thermo Scientific) were washed three times in Co-IP lysis buffer and incubated with 5 μg FLAG-M2 antibody (Sigma) for 1 hour at room temperature with rotation. After incubation, beads were washed three times in lysis buffer and stored on ice. For each immunoprecipitation, one confluent 10 cm plate of HEK293T cells was used. Cells were washed twice with ice-cold PBS, resuspended in Co-IP lysis buffer, and lysed by passing the suspension through a 26G needle. Lysates were centrifuged at 18,000 x g for 5 minutes at 4°C and the supernatant was transferred to a new tube. After removing input controls, the cleared lysate was incubated with the FLAG-M2–bound Dynabeads for 2 hours at 4°C with rotation. Following the incubation, beads were washed three times with low-detergent wash buffer and once with detergent-free buffer using a magnetic stand. During the final wash, beads were transferred to a new tube. For elution, beads were resuspended in 75 μl 2X Laemmli buffer and incubated at 70°C for 5 minutes with vigorous shaking.

### AlphaFold structure predictions

Alphafold3 predictions (*72*) were generated for EEF1G (Uniprot ID P26641) in complex with either oxidized glutathione (CDS code: GDS) or reduced glutathione (CDS code: GSH). For each model, the N-terminal GST domain of EEF1G (residues 1-230) was superimposed onto the corresponding region of an EEF1G-EEF1D crystal structure (PDB: 5JPO) to generate models shown in **Fig, 6, H and I**. Alphafold3 predictions were performed using the BioHPC supercomputing facility located in the Lyda Hill Department of Bioinformatics at UT Southwestern Medical Center.

### Mass photometry

Mass photometry measurements were performed using a Refeyn TwoMP instrument. Glass coverslips for measurements were thoroughly washed with Milli-Q water and isopropanol, followed by drying with nitrogen gas. Silicon wells were applied to the washed coverslips for droplet dilution of samples. Following SEC-buffer addition to wells, the instrument was focused using the droplet dilution auto-focus feature. For data acquisition, protein standards or purified protein complexes were rapidly diluted in SEC-buffer by pipetting and 60 second movies of binding events were subsequently recorded for each sample, capturing approximately 10,000 events per recording. Bovine serum albumin (BSA) and gamma globulin (GG) were used as mass standards. Videos were processed and analyzed using the DiscoverMP software (Refeyn).

### Analysis of disome and ZAKα selective ribosome profiling datasets

Publicly-available disome profiling (GSE134541) (*45*) and ZAKα selective ribosome profiling (GSE141459) (*7*) datasets were obtained from Gene Expression Omnibus (GEO) and adapters were trimmed using Cutadapt (v4.9). Reads were first aligned to ribosomal RNAs (rRNAs) (*86*), transfer RNAs (tRNAs) (*87*), and small nucleolar RNAs (snoRNAs) (*88*) using STAR (v2.7.1a) (*89*). Unmapped reads from this step were aligned to the mouse (GRCm38) or human (GRCh38) reference genomes. Genome annotations were obtained from NCBI for mouse (GCF_000001635.26_GRCm38.p6) and human (GCF_000001405.40-RS_2024_08). For each aligned read, the position of the 13th base from the 3′ end of the read was designated as the P-site.

## Statistical analysis

All experiments were repeated with a minimum of three biological replicates to determine statistical significance. One-sample t-test, Student’s t-test, or 1-way and 2-way ANOVA were calculated using GraphPad Prism 10.10.0. Values are reported as mean ± SD in all figures.

## Acknowledgements

We thank John Doench, Xiaojun Lian, Tobias Meyer, David Root, Didier Trono, and Feng Zhang for plasmids; Vanessa Schmid and Jo Wagner in the McDermott Center Next Generation Sequencing Core at UT Southwestern for assistance with high-throughput sequencing; Björn Grüning and Rolf Backofen for access to the Galaxy Europe platform; and Raghu Chivukula, Kathryn O’Donnell, and members of the Mendell laboratory for critical feedback on the manuscript.

## Author contributions

F.R. and J.T.M. designed the experiments and interpreted the results.

F.R. and C.L. conducted experiments. H.Z. performed bioinformatic analysis of ribosome profiling data. F.R. and J.T.M wrote the manuscript.

## Data and materials availability

All reagents generated in this study are available upon request from J.T.M. with a completed Materials Transfer Agreement. High-throughput sequencing data from the CRISPR screen have been deposited in GEO (GSE299990).

## Supplementary Materials

Figs. S1 to S11

Tables S1 to S4

