## Supplementary Material for "Oxidative stress sensing by the translation elongation machinery promotes production of detoxifying selenoproteins"

**The PDF file includes:**

Figs. S1 to S11

Table S4

**Other Supplementary Materials for this manuscript include the following:**

Table S1 to S3 (Excel)

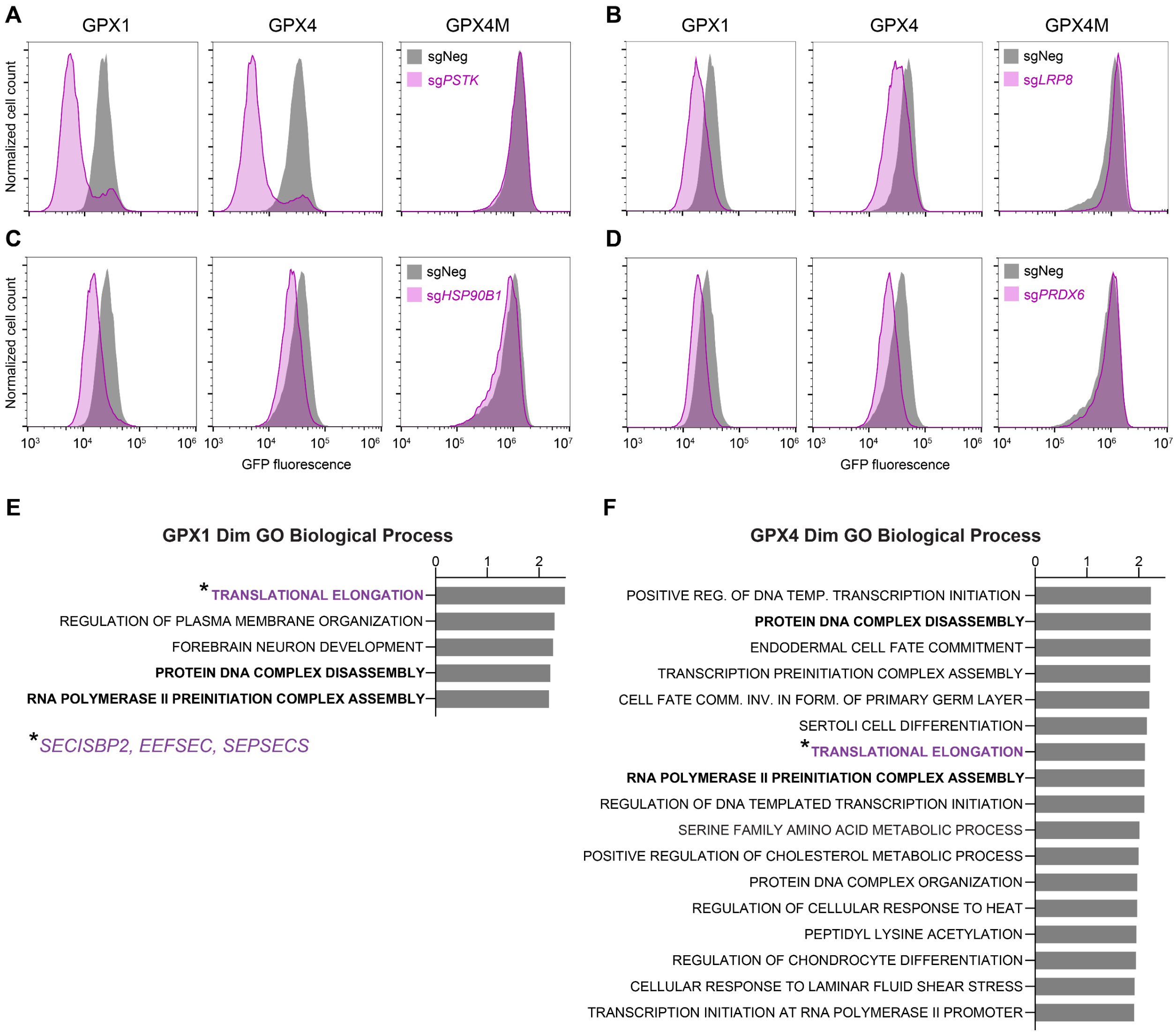

**Figure S1. Validation and analysis of CRISPR screen hits that decreased expression of selenocysteine reporters, related to Fig. 1.** (**A**) to (**D**) Flow cytometry histograms of reporter cell lines transduced with lentiviral constructs encoding Cas9 and the indicated sgRNAs. (**E**) and (**F**) GSEA of dim hits from GPX1 (E) or GPX4 (F) screens. Normalized enrichment scores (NES) of all Gene Ontology (GO) biological process gene sets with false discovery rate (FDR) < 0.25 are shown. 'Translational Elongation' gene set includes core selenocysteine decoding factors SECISBP2, EEFSEC, and SEPSECS (indicated with asterisks). Gene sets labeled in bold text were enriched in both screens and are related to transcription.

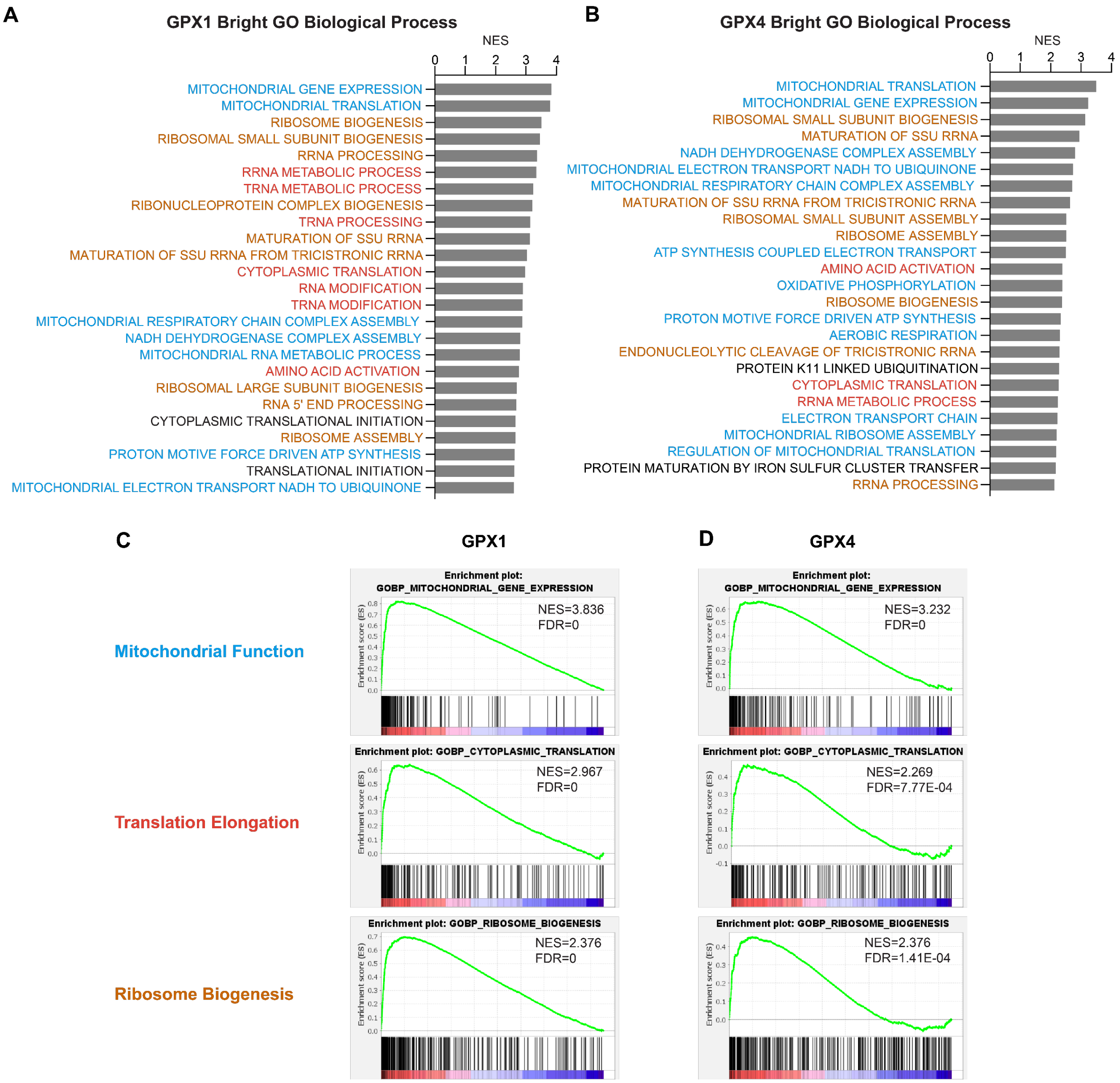

**Figure S2. Analysis of CRISPR screen hits that increased expression of selenocysteine reporters, related to Fig. 2.** (**A**) and (**B**) GSEA of bright hits from GPX1 (A) or GPX4 (B) screens. Normalized enrichment scores (NES) of the top 25 GO biological process gene sets are shown (all FDR < 0.25). Gene sets are categorized into mitochondrial function (blue), translation elongation (red), and ribosome biogenesis (orange). (**C**) and (**D**) GSEA plots of representative gene sets enriched in GPX1 (C) and GPX4 (D) screens.

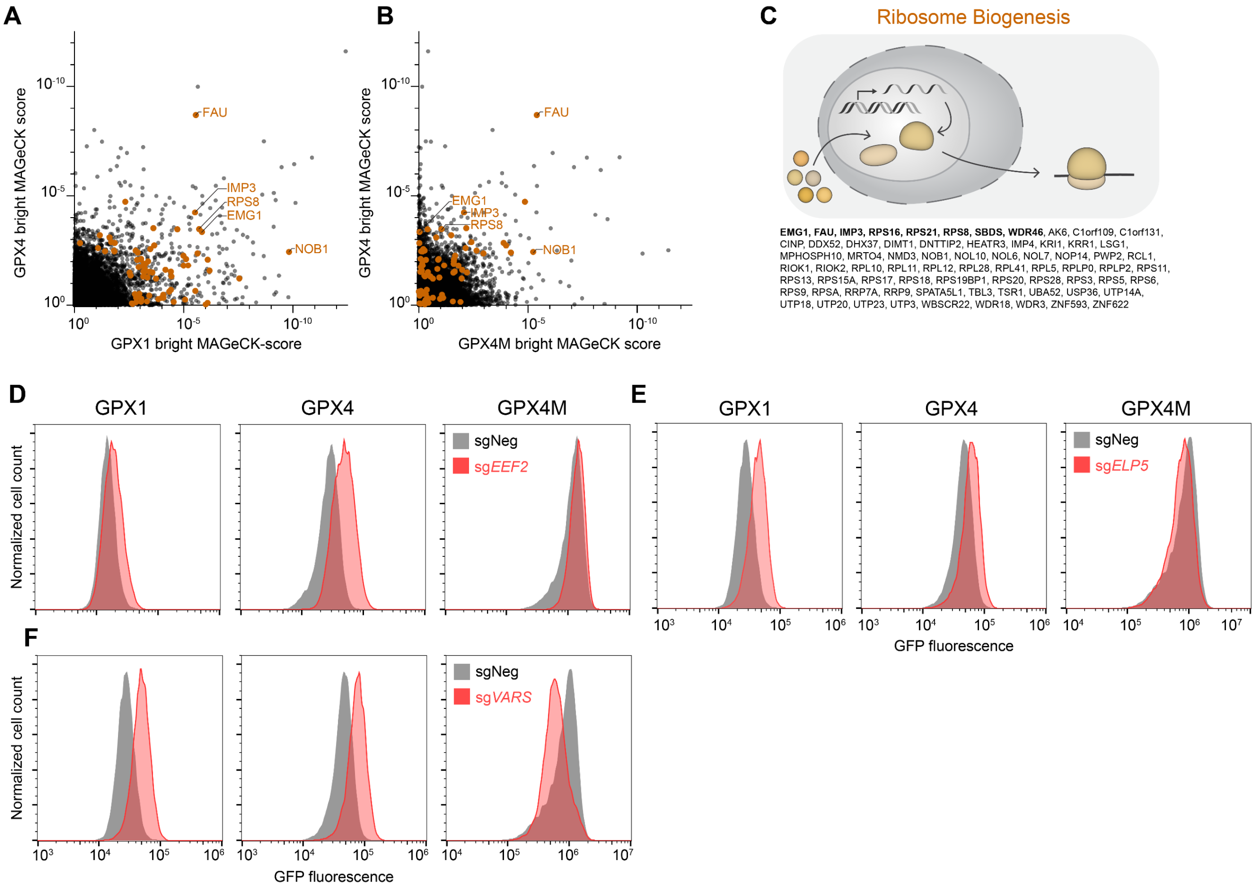

**Figure S3. Analysis and validation of hits related to ribosome biogenesis and translation that increased expression of selenocysteine reporters, related to Figs. 2 and 3.** (**A**) and (**B**) Results of the bright screen for negative regulators of selenocysteine decoding. MAGeCK gene enrichment scores plotted for GPX4 *vs.* GPX1 reporters (A) or GPX4 *vs.* GPX4M reporters (B). Significantly enriched genes in either the GPX1 or GPX4 screen (p<0.01) with roles related to ribosome biogenesis are labeled in orange. (**C**) Significantly enriched genes in either the GPX1 or GPX4 bright screen (p<0.01) with functions related to ribosome biogenesis. Significant hits in both screens are labeled in bold. (**D**) to (**F**) Flow cytometry histograms of GPX reporter cell lines transduced with lentiviral constructs encoding Cas9 and the indicated sgRNAs.

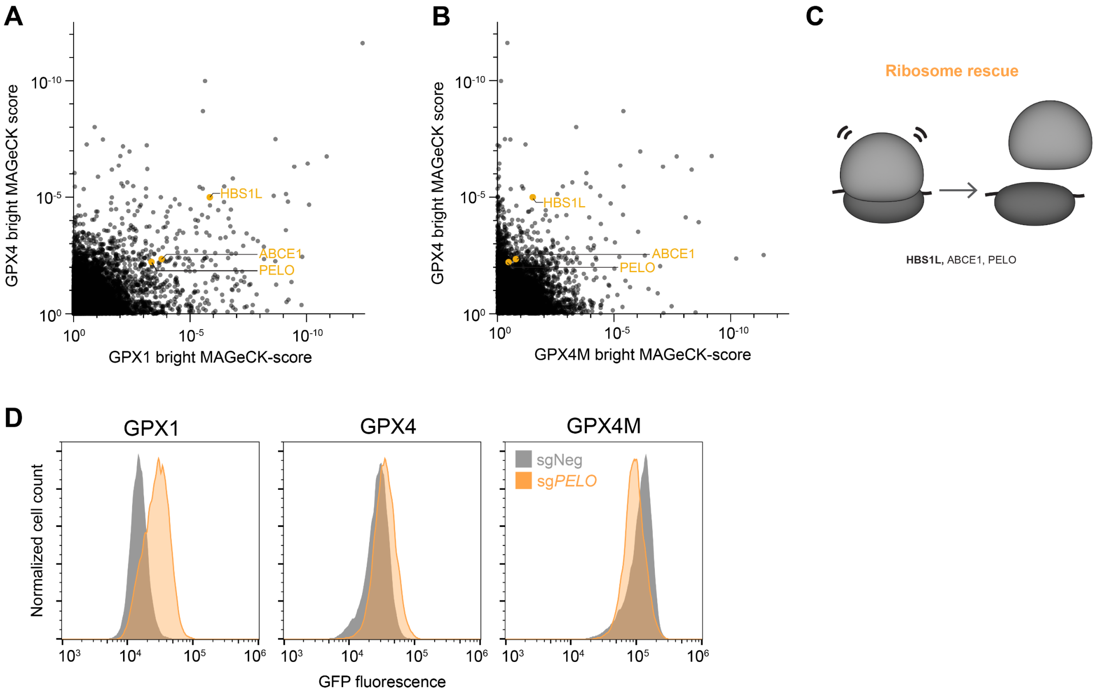

**Figure S4. Loss of ribosome rescue factors increases selenocysteine decoding, related to Fig. 3.** (**A**) and (**B**) Results of the bright screen for negative regulators of selenocysteine decoding. MAGeCK gene enrichment scores plotted for GPX4 *vs.* GPX1 reporters (A) or GPX4 *vs.* GPX4M reporters (B). Significantly enriched genes in either the GPX1 or GPX4 screen ($p<0.01$) with roles related to ribosome rescue are labeled in yellow. (**C**) Significantly enriched genes in either the GPX1 or GPX4 bright screen ($p<0.01$) with functions related to ribosome rescue. HBS1L (bold) was a hit in both screens. (**D**) Flow cytometry histograms of GPX reporter cell lines transduced with lentiviral constructs encoding Cas9 and the indicated sgRNAs.

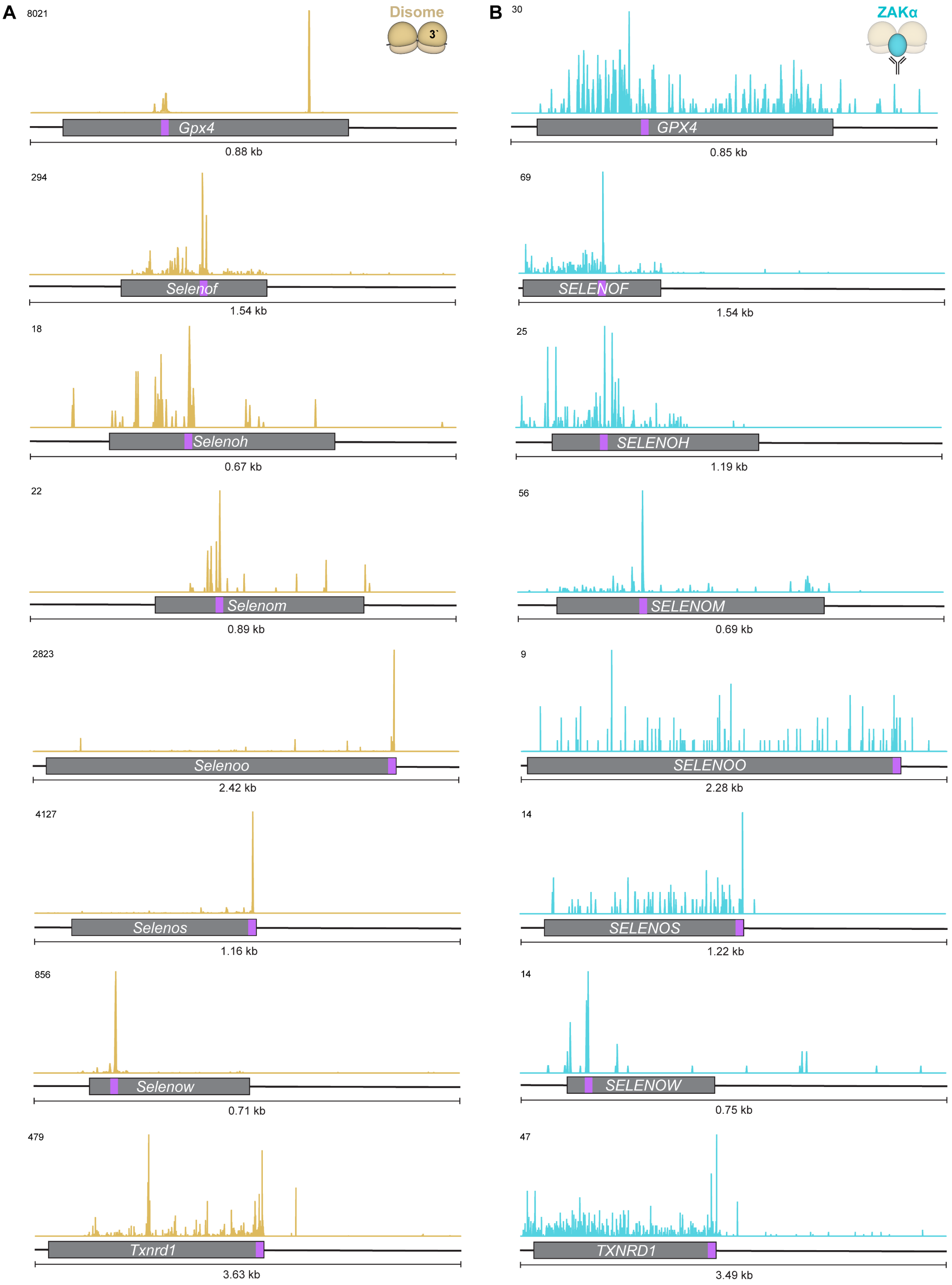

A
Disome
3`
8021
Gpx4
0.88 kb
294
Selenof
1.54 kb
18
Selenoh
0.67 kb
22
Selenom
0.89 kb
2823
Selenoo
2.42 kb
4127
Selenos
1.16 kb
856
Selenow
0.71 kb
479
Txnrd1
3.63 kb
B
ZAKα
30
GPX4
0.85 kb
69
SELENOF
1.54 kb
25
SELENOH
1.19 kb
56
SELENOM
0.69 kb
9
SELENOO
2.28 kb
14
SELENOS
1.22 kb
14
SELENOW
0.75 kb
47
TXNRD1
3.49 kb

**Figure S5. Ribosome collisions frequently occur at selenocysteine codons, related to Fig. 3.** (**A**) and (**B**) Disome profiling (A) and ZAKα selective ribosome profiling (B) read coverage on selenoprotein-encoding transcripts (*1, 2*). For disomes, reads were plotted based on position of the P-site of the leading ribosome. Coding sequences indicated with grey boxes, selenocysteine codons indicated in purple. Numbers in upper left of each plot indicate scale of coverage plots (read counts).

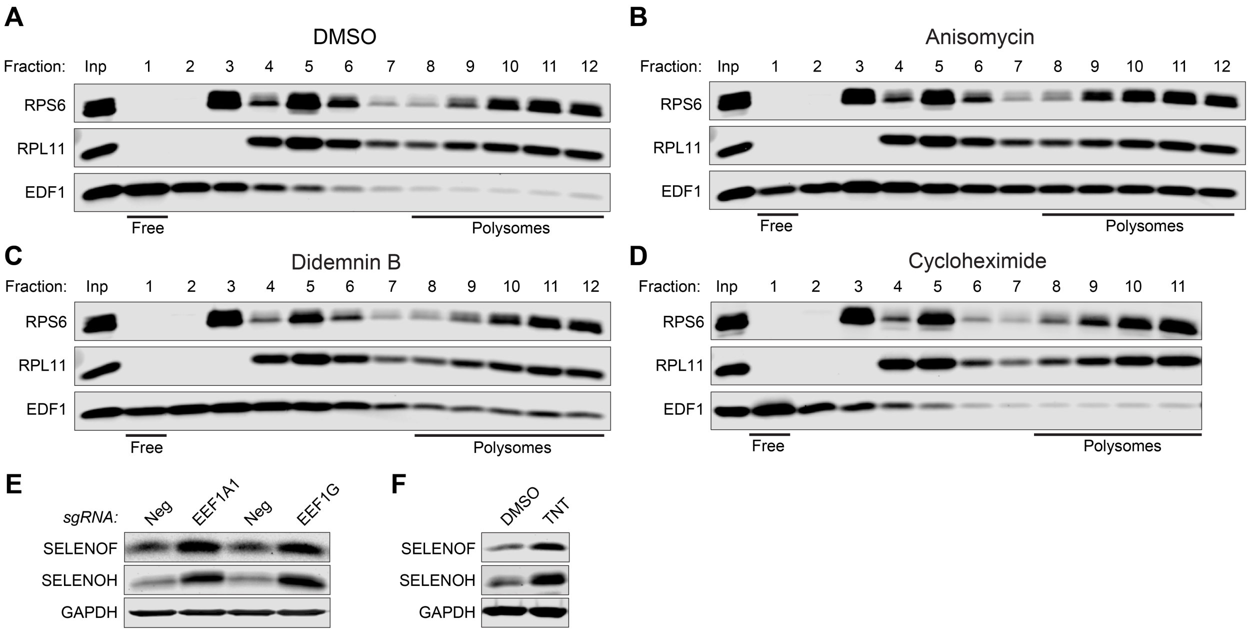

**Figure S6. Effects of translation elongation inhibitors on ribosome collisions and selenoprotein levels, related to Fig. 3.** (**A**) to (**D**) Western blot analysis of sucrose gradient fractions after treatment with DMSO (A), 0.5 μg/ml anisomycin (B), 0.5 μM didemnin B (C), or 0.5 μg/ml cycloheximide (D) for 7 minutes. Inp, input prior to sedimentation. (**E**) and (**F**) Western blot analysis of HCT116 cells following lentiviral delivery of Cas9 and the indicated sgRNAs (E) or after treatment with DMSO or 0.5 μM TNT (F).

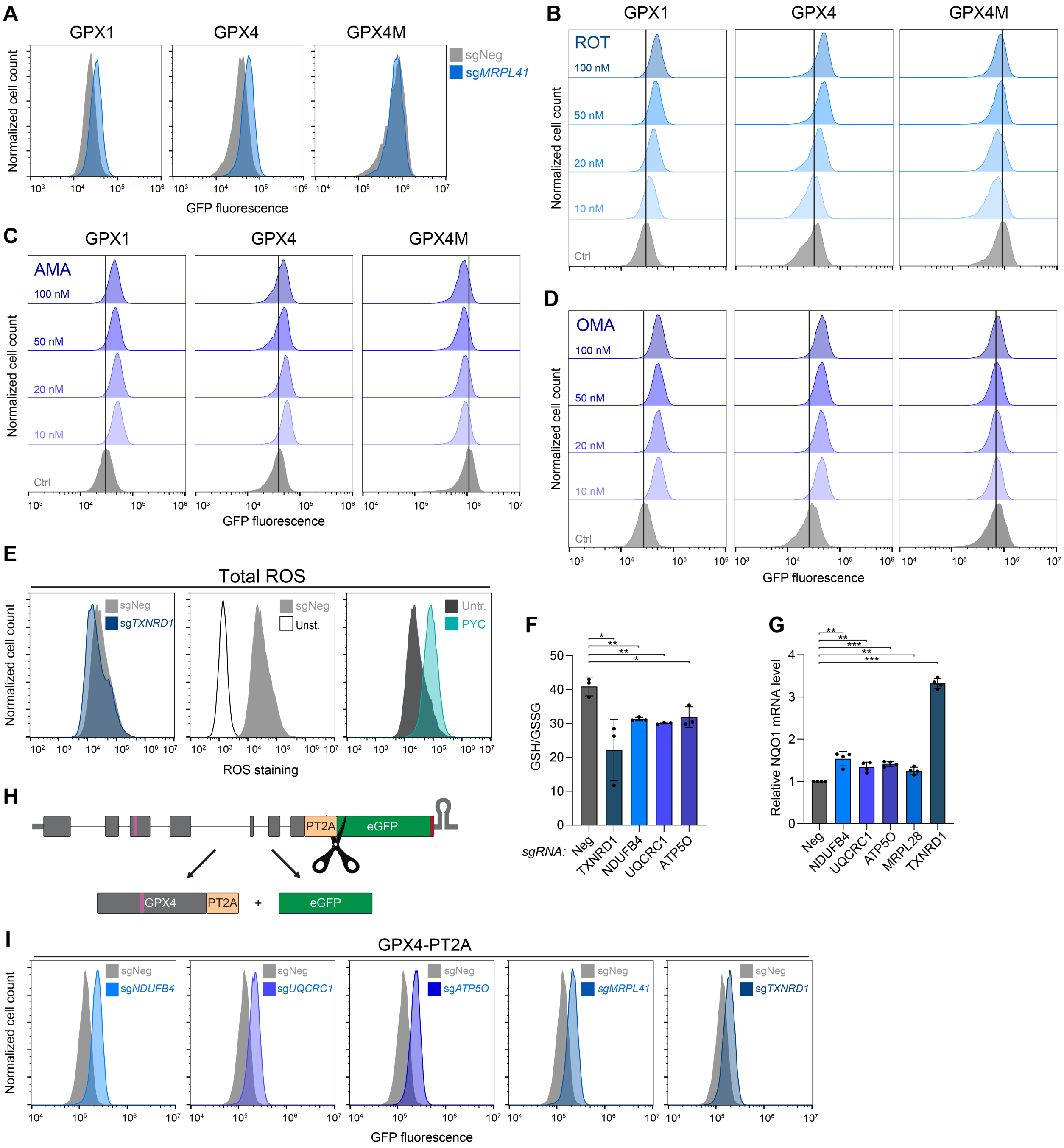

**Figure S7. Oxidative stress and redox imbalance increase selenocysteine decoding, related to Fig. 4.** (**A**) Flow cytometry histograms of GPX reporter cell lines transduced with lentiviral constructs encoding Cas9 and the indicated sgRNAs. (**B**) to (**D**) Flow cytometry histograms of GPX reporter cell lines treated with increasing doses of electron transport chain inhibitors for 72 hours. ROT, rotenone (B); AMA, antimycin A (C); OMA, oligomycin A (D). (**E**) Total cellular ROS quantification by flow cytometry in HCT116 cells. Left: Cells were transduced with Cas9 and the indicated sgRNAs. Middle: Cells transduced with Cas9 and sgNeg were unstained or stained for ROS. Right: Cells with or without 30 minutes of treatment with 200 μM of the ROS-inducing agent pyocyanin (PYC) as a positive control. (**F**) Luminescence-based quantification of GSH/GSSG ratios in HCT116 cells following transduction with lentiviral constructs encoding Cas9 and the indicated sgRNAs. Mean ± SD plotted. n=3 biological replicates. *P* values were calculated by unpaired student's t-test. *p < 0.05, **p < 0.01. (**G**) qRT-PCR analysis of *NQO1* transcript abundance in HCT116 cells transduced with lentiviral constructs expressing Cas9 and the indicated sgRNAs. Expression was normalized to housekeeping gene *Oaz1* and negative control sgRNA. Mean ± SD plotted. n=4 biological replicates. *P* values were calculated by one-sample t-test. **p < 0.01, ***p < 0.001. (**H**) Schematic of GPX4-PT2A-eGFP reporter. A self-cleavable PT2A peptide was inserted between eGFP and GPX4 to rule out post-translational regulation of GPX4 protein stability. (**I**) Flow cytometry histograms of GPX4-PT2A-eGFP reporter cells transduced with lentiviral constructs encoding Cas9 and the indicated sgRNAs.

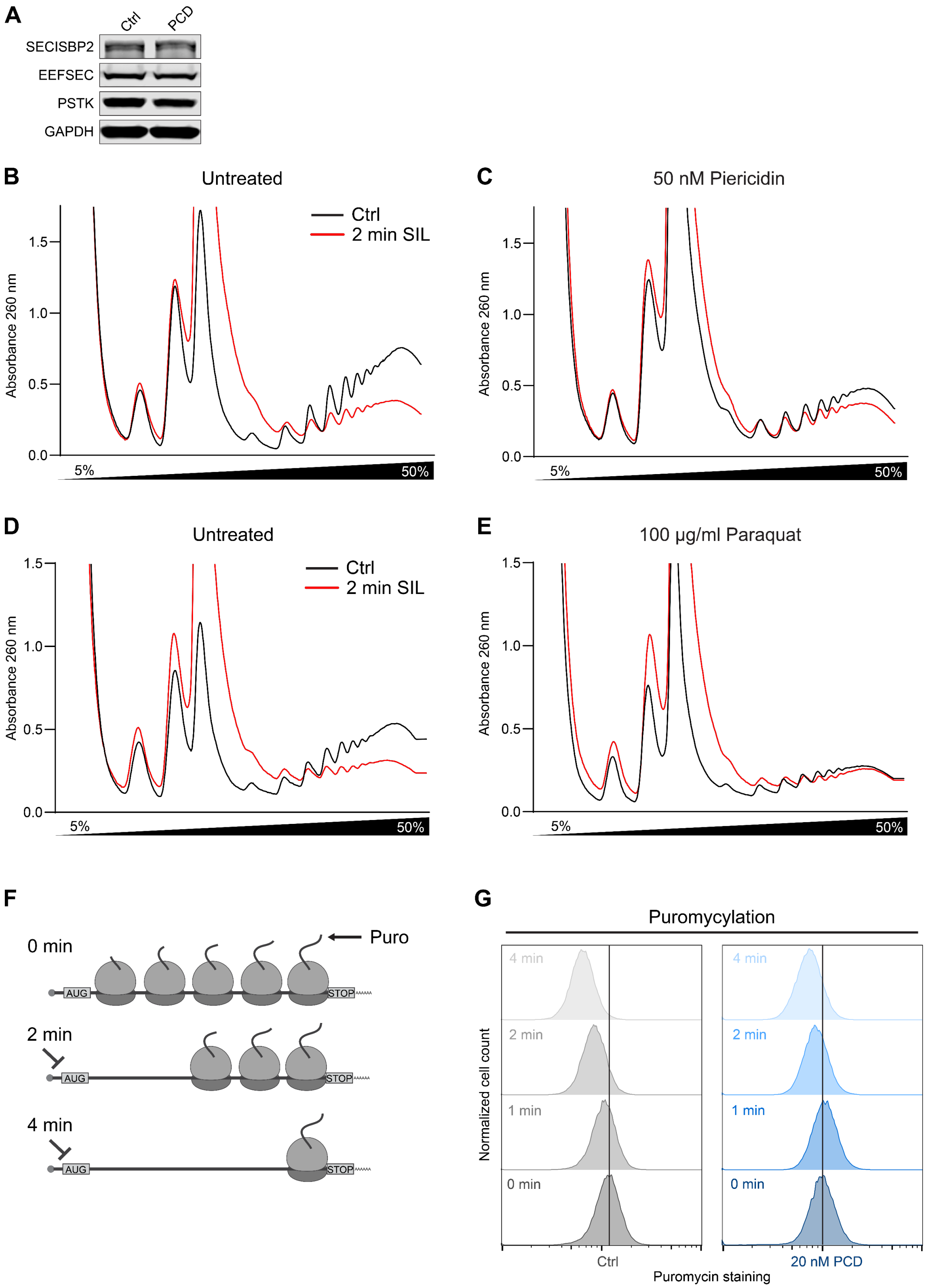

**Figure S8. Oxidative stress slows translation elongation in mammalian cells, related to Fig. 5.** (**A**) Western blot analysis of core selenocysteine decoding factors in HCT116 cells after treatment with 20 nM piericidin (PCD) for 72 hours. (**B**) to (**E**) Ribosome runoff experiments in HCT116 cells with or without 18 hours of treatment with 50 nM PCD (B and C) or 100 µg/ml paraquat (D and E). Polysome abundance was monitored by sucrose gradient sedimentation before and after 2 minutes of treatment with the translation initiation inhibitor silvestrol (SIL; 2

μM). (**F**) Schematic of puromycylation ribosome runoff assays. Cells were treated for 0 to 4 minutes with silvestrol to block initiation and induce ribosome runoff. After puromycin addition, cells were fixed and stained for puromycin incorporation to quantify nascent peptides remaining. (**G**) Flow cytometry analysis of puromycylation ribosome runoff assays. Untreated HCT116 cells (Ctrl) or cells treated for 18 hours with 20 nM PCD were analyzed.

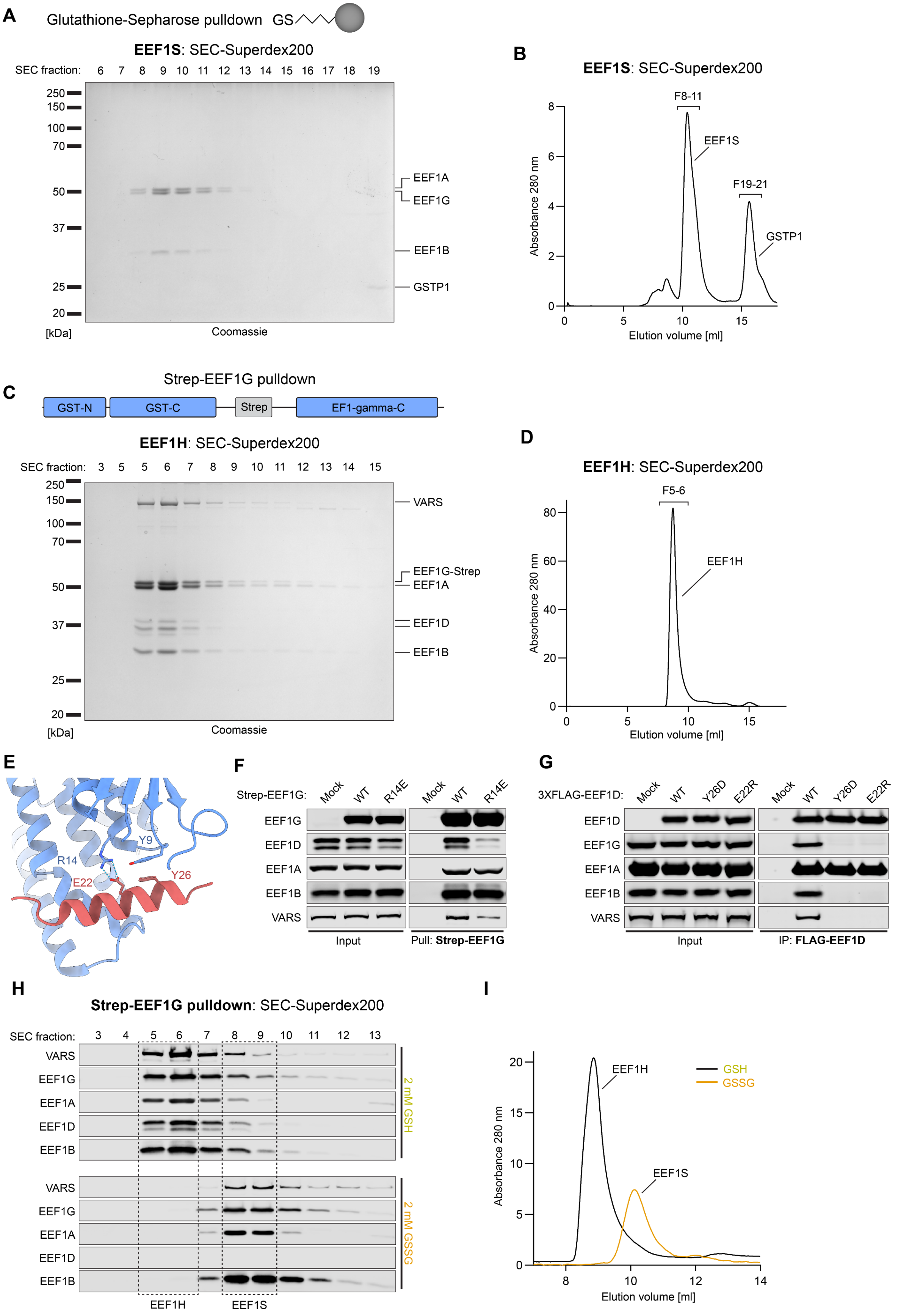
A
Glutathione-Sepharose pulldown GS
EEF1S: SEC-Superdex200
SEC fraction: 6 7 8 9 10 11 12 13 14 15 16 17 18 19
250
150
100
70
50
37
25
20
[kDa]
Coomassie
EEF1A
EEF1G
EEF1B
GSTP1
B
EEF1S: SEC-Superdex200
F8-11
EEF1S
F19-21
GSTP1
Absorbance 280 nm
Elution volume [ml]
C
Strep-EEF1G pulldown
GST-N
GST-C
Strep
EF1-gamma-C
EEF1H: SEC-Superdex200
SEC fraction: 3 5 5 6 7 8 9 10 11 12 13 14 15
VARS
EEF1G-Strep
EEF1A
EEF1D
EEF1B
[kDa]
Coomassie
D
EEF1H: SEC-Superdex200
F5-6
EEF1H
Absorbance 280 nm
Elution volume [ml]
E
Y9
R14
E22
Y26
F
Strep-EEF1G: Mock WT R14E Mock WT R14E
EEF1G
EEF1D
EEF1A
EEF1B
VARS
Input
Pull: Strep-EEF1G
G
3XFLAG-EEF1D: Mock WT Y26D E22R Mock WT Y26D E22R
EEF1D
EEF1G
EEF1A
EEF1B
VARS
Input
IP: FLAG-EEF1D
H
Strep-EEF1G pulldown: SEC-Superdex200
SEC fraction: 3 4 5 6 7 8 9 10 11 12 13
VARS
EEF1G
EEF1A
EEF1D
EEF1B
2 mM GSH
2 mM GSSG
EEF1H
EEF1S
I
EEF1H
EEF1S
GSH
GSSG
Absorbance 280 nm
Elution volume [ml]

**Figure S9. Characterization EEF1 complexes and the effect of glutathione binding on their composition, related to Fig. 6.** (**A**) and (**B**) Glutathione-Sepharose pulldown from HEK293T lysates cells followed by Superdex200 size exclusion chromatography (SEC). SDS-PAGE and Coomassie staining of collected fractions (A) and SEC UV absorbance trace at 280 nm (B) shown. (**C**) and (**D**) StreptactinXT pulldown of endogenous strep-tagged EEF1G from HEK293T cells followed by Superdex200 SEC. SDS-PAGE and Coomassie staining of collected fractions (C) and SEC UV absorbance trace at 280 nm (D) shown. (**E**) Interaction interface of the EEF1D N-terminal helix (aa 12-32) and the EEF1G GST-N domain (PDB: 5JPO). Key residues predicted to be important for the interaction are indicated. (**F**) and (**G**) Mutagenesis of residues that mediate the EEF1G:EEF1D interaction. Western blot analysis following pulldown of wild type or mutant strep-EEF1G (F) or 3XFLAG-EEF1D (G). (**H**) and (**I**) Western blot analysis of SEC fractions (H) and SEC UV absorbance trace at 280 nm (I) following pulldown of endogenous strep-EEF1G in the presence of reduced (GSH) or oxidized (GSSG) glutathione.

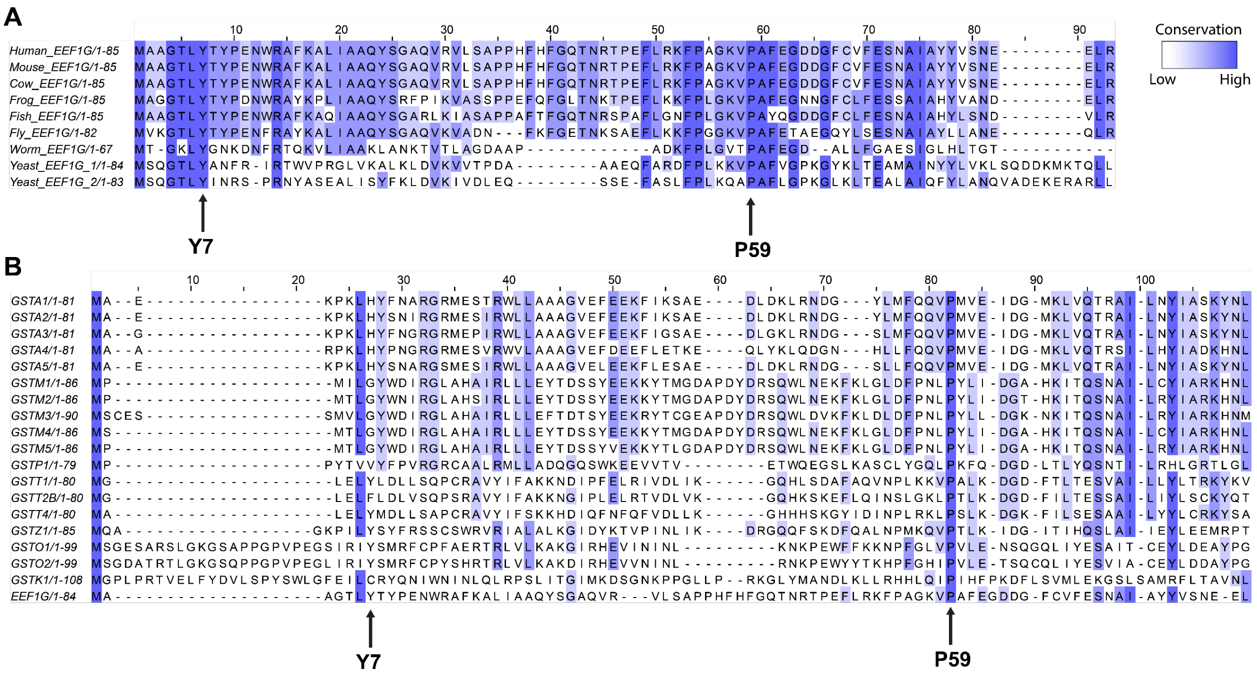

**Figure S10. Conservation of the EEF1G GST-N domain and similarity to canonical GSTs, related to Fig. 7.** (**A**) and (**B**) EEF1G GST-N protein alignments. Constraint-based Multiple Alignment Tool (COBALT) protein alignments of human EEF1G with orthologs from other eukaryotes (A) or the GST-N domains of all annotated canonical human glutathione transferases (B).

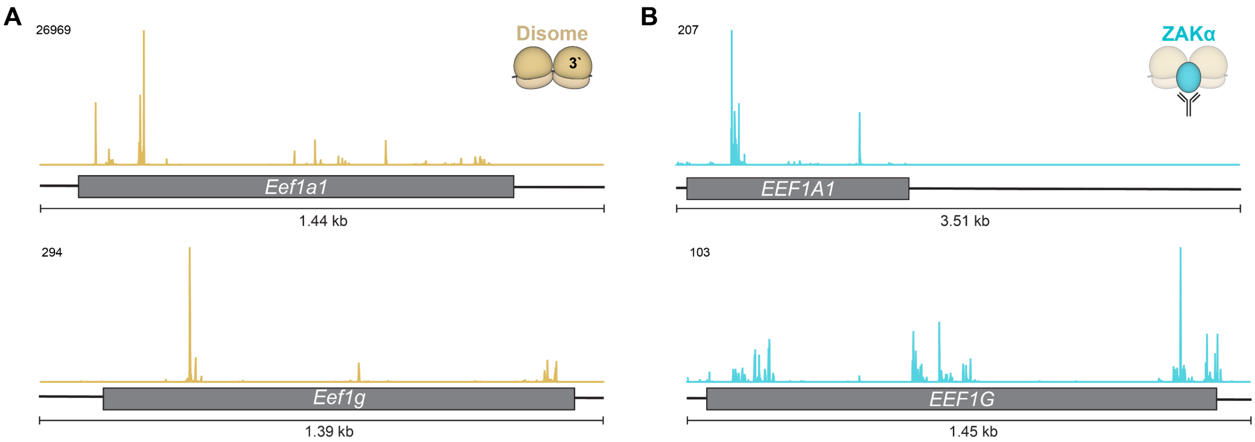

**Figure S11. Ribosome collisions on transcripts encoding translation elongation factors, related to Discussion.** (**A**) and (**B**) Disome profiling (A) and ZAKα selective ribosome profiling (B) read coverage on *EEF1A1* and *EEF1G* (*45, 7)*. For disomes, reads were plotted based on position of the P-site of the leading ribosome. Coding sequences are indicated with grey boxes.

**Table S4. Antibodies and Reagents**

| Antibody | Company | Catalog number |
|---|---|---|
| anti-EDF1 | Fortis Life Science | Cat# A304-039A; RRID:AB_2621288 |
| anti-EEF1A | Proteintech | Cat# 11402-1-AP; RRID:AB_2096966 |
| anti-EEF1B | Proteintech | Cat# 10095-2-AP; RRID:AB_2096987 |
| anti-EEF1D | Santa Cruz | Cat# sc-393731 |
| anti-EEF1G | Santa Cruz | Cat# sc-101035; RRID:AB_2097153 |
| anti-EEF2 | Santa Cruz | Cat# sc-166415; RRID:AB_2277758 |
| anti-EEF2K | Cell Signaling | Cat# 3692; RRID:AB_10694413 |
| anti-EEFSEC | Santa Cruz | Cat# sc-365707; RRID:AB_10844837 |
| anti-FLAG-M2 | Sigma | Cat# F1804; RRID:AB_262044 |
| anti-GAPDH | Santa Cruz | Cat# sc-32233; RRID:AB_627679 |
| anti-GPX1 | Cell Signaling | Cat# 3286; RRID:AB_2067572 |
| anti-GPX4 | Proteintech | Cat# 67763-1-Ig; RRID:AB_2909469 |
| anti-GSTP1 | Santa Cruz | Cat# sc-66000; RRID:AB_2279559 |
| anti-p38 | Santa Cruz | Cat# sc-7972; RRID:AB_628079 |
| anti-phospho-EEF2 | Cell Signaling | Cat# 2331; RRID:AB_10015204 |
| anti-phospho-p38 | Cell Signaling | Cat# 4511; RRID:AB_2139682 |
| anti-PSTK | Invitrogen | Cat# PA5-113118; RRID:AB_2867852 |
| anti-Puromycin-Alexa488 | Sigma | Cat# MABE343-AF488; RRID:AB_2736875 |
| anti-RPL11 | Proteintech | Cat# 16277-1-AP; RRID:AB_2181292 |
| anti-RPS6 | Fortis Life Science | Cat# A300-556A; RRID:AB_477987 |
| anti-SECISBP2 | Proteintech | Cat# 12798-1-AP; RRID:AB_2186555 |
| anti-SELENOF | Aviva Biosystems | Cat# ARP48248_P050 |
| anti-SELENOH | Sigma | Cat# HPA048362; RRID:AB_2680370 |
| anti-TXNRD1 | Santa Cruz | Cat# sc-28321; RRID:AB_628405 |

| | | |
|---|---|---|
| anti-VARS | Proteintech | Cat# 67935-1-Ig;<br>RRID:AB_2918687 |
| anti-β-Actin | Santa Cruz | Cat# sc-69879;<br>RRID:AB_1119529 |
| anti-Mouse secondary IR Dye 800CW | LI-COR | Cat# 926-32212;<br>RRID:AB_621847 |
| anti-Mouse secondary IR Dye 680RD | LI-COR | Cat# 926-68072;<br>RRID:AB_10953628 |
| anti-Rabbit secondary IR Dye 800CW | LI-COR | Cat# 926-32213;<br>RRID:AB_621848 |
| anti-Rabbit secondary IR Dye 680RD | LI-COR | Cat# 926-68073;<br>RRID:AB_10954442 |
| **Reagent** | **Company** | **Catalog number** |
| Agencourt AMPure XP beads | Beckman Coulter Life Sciences | A63882 |
| Alt-R-electroporation enhancer | IDT | 1075915 |
| Alt-R-HDR enhancer V2 | IDT | 10007910 |
| Alt-R-Spy-Cas9 nuclease V3 | IDT | 1081058 |
| Anisomycin | Cayman Chemicals | 11308 |
| Antimycin A | Sigma | A8674-25MG |
| Cycloheximide | Sigma | C1988 |
| Didemnin B | Cayman Chemicals | 34063 |
| Dulbecco′s Modified Eagle′s Medium (DMEM) | Corning | 10-013-CV |
| Fetal Bovine Serum (FBS) | Sigma | F2442 |
| FuGENE HD transfection reagent | Promega | E2312 |
| Glutathione Sepharose 4B | Sigma | GE17-0756-01 |
| GSH/GSSG-Glo Assay | Promega | V6611 |
| Herculase II Fusion DNA Polymerase | Agilent | 600679 |
| L-Glutathione oxidized | Sigma | G4376-1G |
| L-Glutathione reduced | Sigma | G6529-5G |
| MasterPure Complete DNA and RNA Purification Kit | Lucigen | MC85200 |
| NEBuilder HiFi DNA Assembly Master Mix | New England Biolabs | E2621S |
| Oligomycin A | Sigma | 495455-10mg |
| Paraquat dichloride hydrate | Sigma | 36541-100MG |
| Penicillin-Streptomycin | Sigma | P0781-100ML |
| Piericidin A | Cayman Chemicals | 15379 |
| Polybrene | Millipore | TR-1003-G |
| Power SYBRGreen PCR Master Mix | ThermoFisher Scientific | 4367659 |
| PrimeScript RT Master Mix (Perfect Real Time) | Takara | RR036B |

| | | |
|---|---|---|
| PrimeSTAR GXL DNA Polymerase | Takara | R050A |
| Protease Inhibitor Cocktail Set III, EDTA-Free | Millipore | 539134 |
| Puromycin | Invivogen | ant-pr-1 |
| Qubit dsDNA HS Assay Kit | ThermoFisher Scientific | Q32851 |
| RNasin Ribonuclease Inhibitors, recombinant | Promega | N2515 |
| ROS-ID Total ROS detection kit | Enzo | ENZ-51011 |
| Rotenone | Sigma | R8875-1G |
| Silvestrol | MedChem Express | HY-13251 |
| Strep-Tactin XT Sepharose | Cytiva | 29401324 |
| Ternatin | Cayman Chemicals | 20691 |
